# The TET-dependent DNA demethylation pathway is the driving force of hematopoiesis

**DOI:** 10.64898/2026.04.29.721744

**Authors:** Galina Dvoriantchikova, Michelle Fleishaker, Henry Barreras, Robert B. Levy, Dmitry Ivanov

**Affiliations:** Bascom Palmer Eye Institute, Department of Ophthalmology, University of Miami Miller School of Medicine, Miami, FL 33136, USA; Illinois Eye and Ear Infirmary, Department of Ophthalmology, University of Illinois College of Medicine, Chicago, IL, 60612, USA; Department of Microbiology and Immunology, University of Miami Miller School of Medicine, Miami, FL, 33136, USA

**Keywords:** TET dioxygenases, DNA demethylation, hematopoiesis, HSPCs, erythroblasts, T and B cells

## Abstract

A complex network of signaling cascades regulates the differentiation of hematopoietic stem and progenitor cells (HSPCs) into mature blood cells during hematopoiesis. We demonstrated that DNA demethylation driven by Ten-Eleven Translocation (TET) enzymes controls the activity of multiple of these signaling cascades. This occurs because dozens of genes, essential for the specification (e.g., Gata1, Tcf7, Bcl11b, Pou2af1) and maturation (e.g., TCR and BCR signaling) of erythroblasts, T cells, and B cells, are highly methylated in HSPCs and must be demethylated by TET enzymes to be activated in their respective blood lineages. Many genes required for the maturation, but not for the specification, of monocytes and granulocytes are highly methylated in HSPCs, a condition that does not impede the emergence of these cells but can significantly impair their function. Loss of TET activity leads to severe impairment of hematopoiesis and results in hematological malignancies.

## Introduction

Hematopoiesis is a complex and critical process for the body’s vital functions that leads to the generation of various blood cell types from hematopoietic stem cells (HSCs)^1–3^. HSCs differentiate into multipotent progenitors (MPPs) that possess the same potential to generate all types of blood cells as HSCs (therefore, these types of cells are grouped together as hematopoietic stem and progenitor cells [HSPCs]) ^1–3^. MPPs further differentiate into committed progenitors with the ability to differentiate only into lymphoid (common lymphoid progenitors or CLPs) or myeloid and erythroid (common myeloid progenitors or CMPs) lineages^1–3^. CLPs differentiate into lymphocytes, including B cells and T cells^1–3^. CMPs undergo further specification, differentiating into megakaryocyte–erythroid progenitors (MEPs; that differentiate into megakaryocytes and erythroblasts/erythrocytes) and granulocyte-monocyte progenitors (GMPs; that differentiate into granulocytes [e.g., neutrophils] and monocytes/macrophages)^1–3^. All these stages of hematopoiesis are tightly controlled by a number of signaling cascades^1–3^. Impaired activity of these signaling cascades leads to blood diseases, many of which can be fatal^1–3^. In this study, we demonstrate for the first time that the activity of many of these cascades is under the control of a single epigenetic pathway that is responsible for DNA demethylation.

Genome-wide DNA methylation patterns are formed by the opposing activities of two gene families: DNA methyltransferases (DNMTs) are responsible for the methylation of cytosines in DNA, while Ten-Eleven Translocation (TET; Tet1, Tet2, and Tet3) dioxygenases are responsible for their demethylation^4,5^. High levels of methylation (hypermethylation) of the promoter and first exon often suppress gene expression^6,7^. Thus, DNA methylation patterns play a central role in determining which genes are active and which are repressed in a cell. The process of DNA demethylation plays a distinctive role in hematopoiesis, since mutations in TET genes and in genes whose products affect TET activity can lead to a range of blood diseases, including acute myeloid leukemia (AML)^8,9^. However, how the TET-dependent DNA demethylation pathway is mechanistically involved in hematopoiesis, and how its disruption drives blood diseases, remain poorly understood. In this study, we address these questions through the use of TET-deficient mice.

It is often assumed that promoters and first exons of genes required for the development of various cell types are hypomethylated (i.e., have low methylation levels) in stem cells from which these cell types originate. As stem cells differentiate into a specific cell type, genes required for alternative cell fates become hypermethylated, while only the genes needed for that specific cell type remain hypomethylated. We discovered that the reverse mechanism operates during hematopoiesis. Our findings reveal that many genes required for the development and function of various blood cell types are highly methylated in HSPCs. During the differentiation of HSPCs into a specific blood cell type, only the genes required for that cell type are selectively demethylated by TET enzymes, while genes required for alternative blood cell fates remain hypermethylated. Lack of TET-dependent DNA demethylation results in HSPCs being unable to differentiate into various blood cell types, leading to HSPC accumulation. This results in a deficiency of vital blood cells alongside a buildup of undifferentiated HSPCs, ultimately causing lethality in TET-deficient mice, a phenotype frequently observed in leukemia. These results indicate that the TET-dependent DNA demethylation pathway exerts global control over hematopoiesis by demethylating dozens of key hematopoietic genes.

## Results

### Many genes required for the specification and maturation of various blood cell types are hypermethylated in HSPCs and become demethylated during HSPC differentiation

We used whole genome bisulfide sequencing (WGBS) data to investigate the changes that occur in DNA methylation patterns during the differentiation of mouse HSPCs into myeloid (monocytes/granulocytes and erythroblasts) and lymphoid (T cells and B cells) lineages. To this end, monocytes/granulocytes and erythroblasts were isolated from the bone marrow of wild type mice using anti-CD11b and anti-Ter-119 microbeads, respectively, and the magnetic cell separation strategy. T cells were isolated from the spleen using magnetically labeled antibodies in accordance with the MACS separation protocol. We utilized published WGBS data to analyze the methylation patterns of HSPCs and B cells^10–12^. Comparative analysis of DNA methylation patterns showed that representatives of the myeloid lineage cluster closer to HSPCs than representatives of the lymphoid lineage (**Fig. 1A**). Analysis of differentially methylated cytosines (DMCs) showed that while the number of DMCs with decreasing and increasing levels of methylation during HSPCs differentiation is approximately equal for representatives of the lymphoid lineage, the methylation of most DMCs decreases in representatives of the myeloid lineage (**Fig. 1B**). This was especially noticeable in erythroblasts, in which almost all cytosines were demethylated. Using the methylKit Bioconductor package, in addition to individual cytosines, we identified genomic regions with similar methylation levels in HSPCs, monocytes/granulocytes, erythroblasts, T cells, and B cells. We identified four types of regions, two of which correspond to hypomethylated genomic regions (1 and 2) and two correspond to hypermethylated genomic regions (3 and 4). Among all the types of blood cells examined, erythroblasts stand out in particular, as their genome is the least methylated (**Fig. 1C**). We compiled a list of genes whose promoters and first exons lie within the genomic regions exhibiting statistically significant changes in methylation levels in the pairs: Monocyte/Granulocytes vs. HSPCs, Erythroblasts vs. HSPCs, T cells vs. HSPCs, B cells vs. HSPCs (**Fig. 1D**, **Supplementary Data S1**). We assumed that changes in the methylation levels of these regions reflected changes in the methylation levels of these genes. While we were unable to identify genes important for hematopoiesis whose methylation increased during HSPC differentiation, we identified numerous genes critical for erythroblast, T cell, and B cell specification and maturation/function whose methylation decreased during HSPC differentiation (**Fig. 1D**, **Supplementary Data S1**). Of particular note are the genes critical for T cell receptor (TCR) signaling (e.g., Cd3d, Cd3g, Cd3e, Sit1, Trat1, Themis), B cell receptor (BCR) signaling (e.g., Cd79a, Cd19, Vpreb3, Ms4a1, Cd52), and for the development of B cells (Pou2af1 and Spib) and erythroblasts (e.g., Gata1, Spta1, Alas2, Tspo2, Cd36) (**Fig. 1D, Supplementary Data S1**). We also found genes that are demethylated during the differentiation of HSPCs into monocytes/granulocytes (**Fig. 1D, Supplementary Data S1**). The products of these genes may affect the maturation/function of these cells but are probably not important for their specification. It should be noted that many genes important for the development and function of various blood cell types are located in hypermethylated genomic regions of HSPCs (**Fig. 1D**). However, when HSPCs differentiate into a specific blood cell type, only the genomic regions containing the genes needed for that blood cell type are demethylated (**Fig. 1D**). Thus, the process of DNA demethylation is blood cell type specific. The exception is erythroblasts, in whose genome many genomic regions are demethylated. (**Fig. 1D, Supplementary Data S1**). Since the approach we described, based on calculating the average methylation levels of genomic regions using the methylKit Bioconductor package, does not account for changes in the methylation levels of individual cytosines near the gene transcription start site (TSS± 1,000 bp; this region contains the promoter and the first exon), the resulting data are qualitative rather than quantitative in nature. To circumvent these limitations and independently validate our results, we utilized the DMRseq Bioconductor package that is free of these constraints and identifies differentially methylated regions (DMRs) based on the methylation levels of individual cytosines. We found that DMR distribution is very similar to what we observed for DMCs (**Fig. 1B, 1E**). Our results indicate that the TSS± 1,000 bp of almost all identified hematopoiesis genes is located in one of these DMRs (**Fig. 1F, Supplementary Data S2**). We were also able to identify additional genes important for the development of T cells (Tcf7 [TCF-1] and Ets1) and for BCR signaling (e.g., Blnk, Cd22) located in these DMRs (**Supplementary Data S3**).

**Figure 1.**
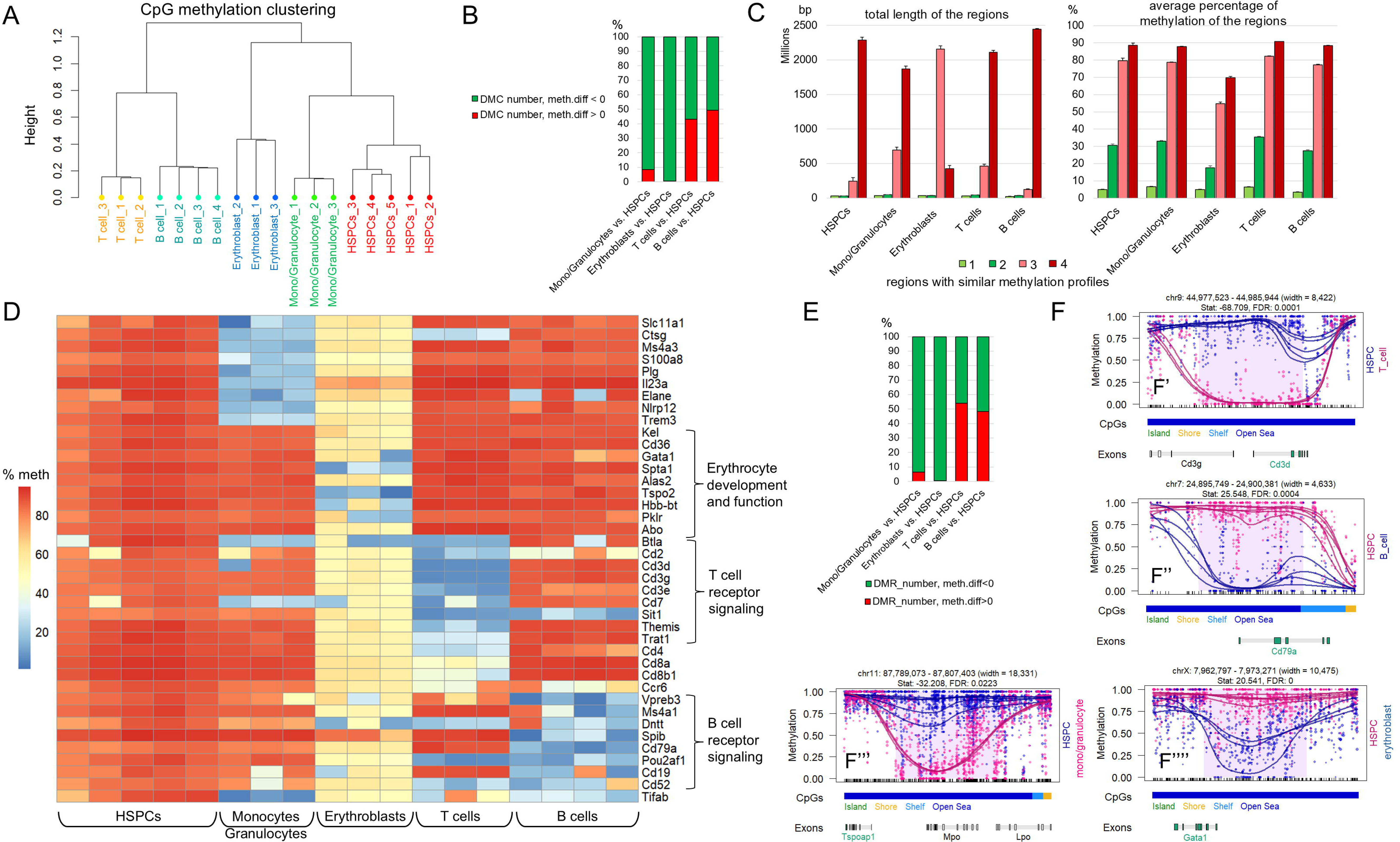
Genes critical for hematopoiesis are hypermethylated in HSPCs and are selectively demethylated during the differentiation of HSPCs into mature blood cells. **A)** Sample clustering based on the similarity of methylation profiles (clustering method: “ward”) was performed using the methylKit Bioconductor package. **B)** Differentially methylated cytosines (DMCs) were identified using the methylKit package (FDR < 0.01). Data on the number of DMCs with reduced (green) and increased (red) methylation relative to HSPCs was used to calculate the percentages in each blood cell type studied. **C)** Four types of genomic regions were identified based on the similarity of their cytosine methylation levels as a result of the segmentation analysis (methylKit). The total length of the regions and their average methylation level were calculated. **D)** Average methylation levels of genomic regions identified in the segmentation analysis and genes critical for hematopoiesis in mice were used to generate the heatmap. **E)** The panel shows the percentages of differentially methylated regions (DMRs) with reduced (green) and increased (red) methylation levels. DMRs with reduced methylation levels are those whose methylation decreases during HSPC differentiation. **F)** The panel provides examples of DMRs in T cells (F’), B cells (F’’), monocytes/granulocytes (F’’’), and erythroblasts (F’’’’) vs. HSPCs. Each subpanel was generated using the DMRseq Bioconductor package.

### DNA demethylation is predominant in human hematopoiesis

To understand whether our findings are species-specific or a general phenomenon, we conducted a similar study, but on human HSPCs, monocytes, erythroblasts, T cells, and B cells. To this end, we analyzed published WGBS data^13–16^. The clustering of cells based on their DNA methylation patterns was similar to what we observed in mice (**Fig. 1A, 2A**). At the same time, compared to mice, demethylation of cytosines (DMC number, meth.diff < 0) during HSPC differentiation was predominant in all lineages (**Fig. 2B**). This was reflected in the fact that the number of genomic regions with reduced methylation levels was high not only in erythroblasts but also in T cells and B cells (**Fig. 2C**). Using the same approach as with the mouse WGBS data, we also compiled a list of genes whose promoter and first exon methylation levels change statistically significantly during hematopoiesis (**Supplementary Data S4**). Analysis of the gene list revealed that many genes important for hematopoiesis are located in hypermethylated regions of HSPCs (**Fig. 2D**, **Supplementary Data S4**). These same genomic regions are demethylated during differentiation of HSPCs into the corresponding blood cell types. One important exception is SPI1 (PU.1), whose methylation is increased in T cells (**Fig. 2D**). We also found that the percentage of DMRs with increased and decreased levels of DNA methylation is consistent with what we observe for DMCs (**Fig. 2B, 2E**). The TSS ±1,000 bp of almost all hematopoietic genes in our list are located in DMRs with reduced methylation levels (**Fig. 2F**, **Supplementary Data S5**). Thus, as in mice, many genes important for the specification and maturation/function of various human blood cell types are hypermethylated in HSPCs (**Fig. 2D**, **Supplementary Data S4**). During HSPC differentiation, only genes important for a specific blood cell type, but not other types of blood cells, are demethylated (**Fig. 2D, 2G**, **Supplementary Data S4, S5**). An exception to this pattern, as in mice, is observed in erythroblasts (**Fig. 1D, 2D**). It should be noted that, as in mice, the genes that are demethylated during the differentiation of HSPCs into monocytes are likely not critical for the specification of these cells (**Fig. 2D**, **Supplementary Data S5**). At the same time, the genes that are demethylated during the differentiation of HSPCs into erythroblasts (e.g., GATA1, KLF1, SPTA1, ALAS2, CD36, HBB), T cells (e.g., CD3D, CD3E, CD3G, TRAT1, SIT1, THEMIS, UBASH3A), and B cells (e.g., CD79A, FCRLA, BLNK, VPREB3) are critical for the specification and maturation/function of these blood cell types (**Fig. 2D**, **Supplementary Data S5, S6**).

**Figure 2.**
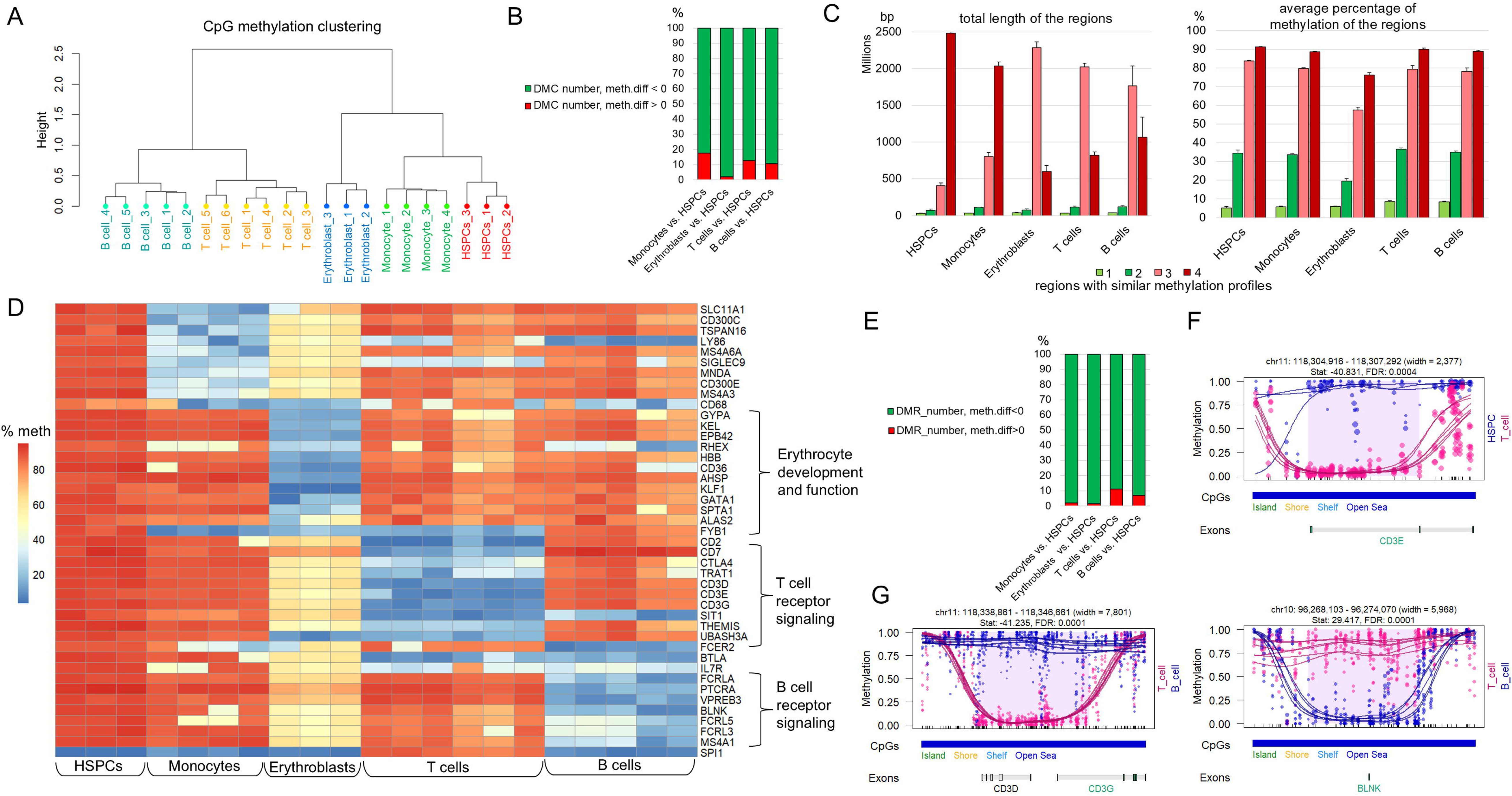
The development of various types of human blood cells is accompanied by demethylation of many genes necessary for their development and function. **A)** Clustering of samples using the methylKit package revealed a close similarity between the methylation profiles of myeloid lineage cells and the methylation profile of HSPCs. **B)** Counting DMCs (FDR < 0.01) in the genomes of human HSPCs vs. blood cell types revealed that those whose methylation reduces during HSPCs differentiation predominate. **C)** The segmentation analysis (methylKit) indicates that among the genomic regions in erythroblasts, T cells, and B cells, those with a lower average level of methylation (compared to HSPCs) predominate. **D)** The heatmap shows the average methylation levels of genomic regions containing promoters and first exons of genes critical for hematopoiesis in humans. **E)** The percentage of DMRs where methylation reduced (green) or increased (red) during human HSPC differentiation. **F)** An example of one of the genes that are demethylated during hematopoiesis and are important for this process, the transcription start site (TSS; ±1,000 bp) of which is located in DMR. **G)** The panel shows examples of genes that are demethylated differently during the differentiation of T cells and B cells from HSPCs.

### Genetic ablation of the TET-dependent DNA demethylation pathway impairs hematopoiesis and leads to leukemia-like pathology

Will HSPCs be able to differentiate into erythroblasts (erythrocytes), T cells, and B cells if the genes responsible for their specification and maturation are not demethylated during hematopoiesis? What would happen to monocytes and granulocytes in this scenario, given that nothing impedes their specification? To answer these questions, we generated mice in which all three Tet1, Tet2, and Tet3 genes responsible for the DNA demethylation process could be deleted by treating them with tamoxifen dissolved in oil (KO, **Fig. 3A**). As a control (C), we used the same animals treated with oil only. In the first month, the experimental tamoxifen-treated (KO, TET-deficient) and control (C) groups did not differ in any way in terms of their behavior or health status (**Fig. 3B**). Differences began to appear in the second month. We observed rapid disease progression in mice treated with tamoxifen (KO). In just one week, it transformed normal-looking animals into very sick ones exhibiting heavy labored breathing. These mice were very pale that was also evidenced upon examination of their femurs and tibiae bones (**Fig. 3C**). In total, this indicated a deficiency of erythrocytes/erythroblasts, which was confirmed by flow cytometry (Cd71/Ter119; **Fig. 3D**). Examination of tissues from TET-deficient (KO) mice also revealed a greatly enlarged (splenomegaly) but pale spleen (**Fig. 3E**). The number of cells in the spleens of KO mice was tenfold greater than the number of cells in the spleens of control (C) animals (**Fig. 3E**). The liver and lungs were also enlarged, which was reflected in their weight (**Fig. 3F**). Analysis of blood cell types in the bone marrow and spleen using flow cytometry showed that the number of T cells (Cd4, Cd8) and B cells (Cd19/B220) was lower, while the number of HSPCs (Cd117), myeloid progenitors (Cd117/Cd11b), and monocytes/granulocytes (Gr-1/Cd11b) was higher in TET-deficient (KO) mice (**Fig. 3G**). We also analyzed the serum of TET-deficient and control mice using liquid chromatography-tandem mass spectrometry (LC-MS/MS). Reduced levels of B cell receptor (BCR) proteins in the serum of TET-deficient vs. control mice were identified (**Fig, 3H**, **Supplementary Data S7**). We also found reduced levels of blood clotting proteins (**Fig, 3H**, **Supplementary Data S7**). These data indicate that the number and/or function of B cells and megakaryocytes/platelets are reduced. Thus, the number of blood cells arising from the differentiation of CLPs and MEPs was reduced, whereas the number of HSPCs and cells derived from GMPs was increased in TET-deficient mice (**Fig. 3I**). Overall, the inactivation of the TET-dependent DNA demethylation pathway hinders the differentiation of HSPCs into CLP and MEP lineages, resulting in severe pathological changes frequently observed in leukemia (e.g., anemia, lymphopenia, low levels of coagulation factors, an elevated HSPC count).

**Figure 3.**
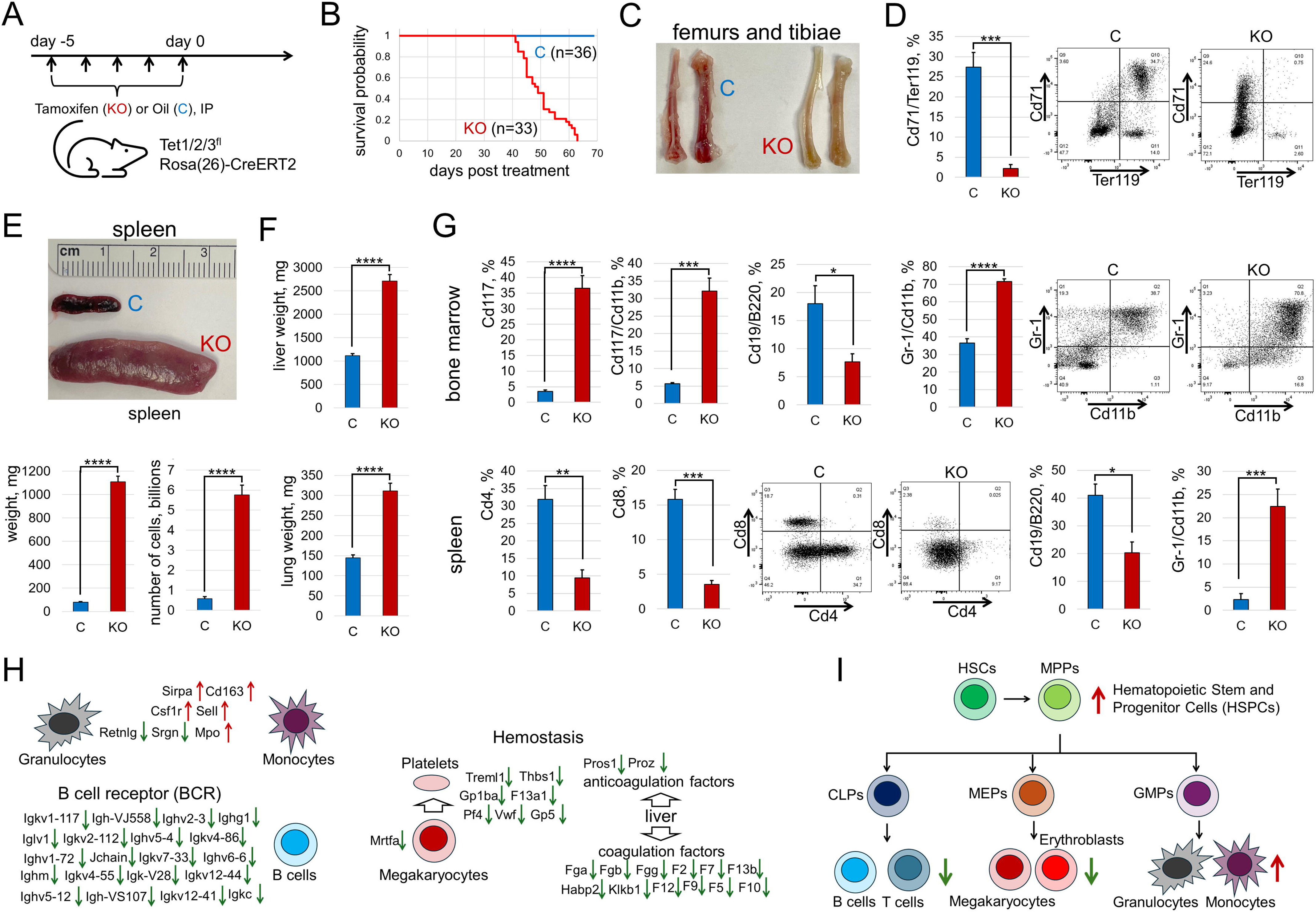
DNA demethylation driven by TET enzymes is essential for hematopoiesis. **A)** Triple TET floxed mice and Rosa(26)-CreERT2 mice were crossed to create mice (heterozygous for CreERT2 and homozygous for TET-floxed) in which all TET genes could be deleted by treating them with tamoxifen. The experimental group (KO) of 2-month-old mice was injected daily with tamoxifen (dissolved in oil) intraperitoneally (IP) for 5 days. The control group of mice (C) with the same genotype was treated only with oil. **B)** Kaplan–Meier curve representing survival probability of C (n = 36) and KO (n = 33) mice was generated after all the tamoxifen-treated (KO) animals had died. Animals were euthanized when they became very sick and it was clear they would die within a few hours. **C)** The representative image of femurs and tibiae show the low erythrocyte content in the tissues of the tamoxifen-treated (KO) mice. **D)** Fluorescence flow cytometry (FFC) was performed to determine the erythroblast/erythrocyte content in the bone marrow of experimental (KO) and control (C) mice (***P < 0.001). **E)** The representative image of experimental (KO) and control (C) spleens, the results of weighing the spleens, and the counting of the number of cells in them all indicate the development of splenomegaly in TET-deficient mice. **F)** The results of weighing the liver and lungs indicate their significant enlargement in tamoxifen-treated (KO) mice. **G)** FFC was performed to analyze hematopoietic cell populations in the spleen and bone marrow of experimental (KO) and control (C) animals (*P < 0.05, **P < 0.01, ***P < 0.001, ****P < 0.0001). **H)** Serum was analyzed using LC-MS/MS. The results indicate that the inactivation of TET enzymes strongly affects the composition of proteins in the blood. **I)** The collected data indicate that the activity of TET enzymes is essential for the differentiation of HSPCs into erythroblasts/erythrocytes, megakaryocytes, T cells, and B cells. The inability of TET-deficient HSPCs to differentiate leads to their accumulation. A green downward arrow indicates a decrease in the number of cells, while a red upward arrow indicates an increase in the number of cells in TET-deficient mice.

### The TET-dependent DNA demethylation pathway controls the expression of many genes critical for T cell specification and maturation/function

To investigate how TET enzyme inactivation affects the development of the lymphoid lineage at the molecular level, we isolated T cells from the spleens of tamoxifen-treated experimental (KO, TET deficient) very sick mice and control (C) mice. The number of isolated T cells from control mice was twice the number of T cells isolated from experimental TET-deficient mice (C: 6.3±1.1×10^6^ vs. KO: 3.2±0.5×10^6^, P-value= 0.0161), which was consistent with our flow cytometry data (**Fig. 3G**). The isolated T cells were used in WGBS and RNA-seq analysis. Using the DMRseq Bioconductor package to analyze our WGBS data, we found that the TSS ±1,000 bp of almost all genes whose promoters and first exons are demethylated during T cell differentiation from HSPCs were located in DMRs with increased methylation (**Fig. 4A**, **Supplementary Data S8**). It should be noted that the isolated cell populations contain both T cells that originated before tamoxifen treatment and those that differentiated from TET-deficient HSPCs after treatment since the lifespan of some T cells can exceed the lifespan of the entire organism. This explains why the methylation level of several genes in TET-deficient (KO) T cells was not very high (**Fig. 4A**, **Supplementary Data S8**). To investigate how high levels of TSS±1,000 bp methylation in TET-deficient T cells affect gene expression, we analyzed our RNA-seq data. We found that the inactivation of TET enzymes had a significant impact on gene expression in T cells (**Fig. 4B**). The expression of almost all identified in our study T cell genes whose TSSs (±1,000 bp) are located in DMRs was reduced (**Fig. 4A**, **4C**, **Supplementary Data S8**). However, in addition to these genes, the expression of at least 70 genes important for T cell specification and maturation/function was also reduced in TET-deficient T cells (**Fig. 4C**, **Supplementary Data S8**). Further analysis revealed that TSSs (±1,000 bp) of many of these genes are located in DMRs with increased methylation (**Supplementary Data S8**). Notably the expression of Cd34 and Kit (Cd117), genes characteristic of HSPCs, was increased (**Fig. 4C**). Overall, we observed reduced expression of genes required for T cell specification (Tcf7 [TCF-1], Bcl11b) and maturation (e.g., TCR signaling genes), which should lead to impaired development and low T cell counts in TET-deficient mice (**Fig. 4D**).

**Figure 4.**
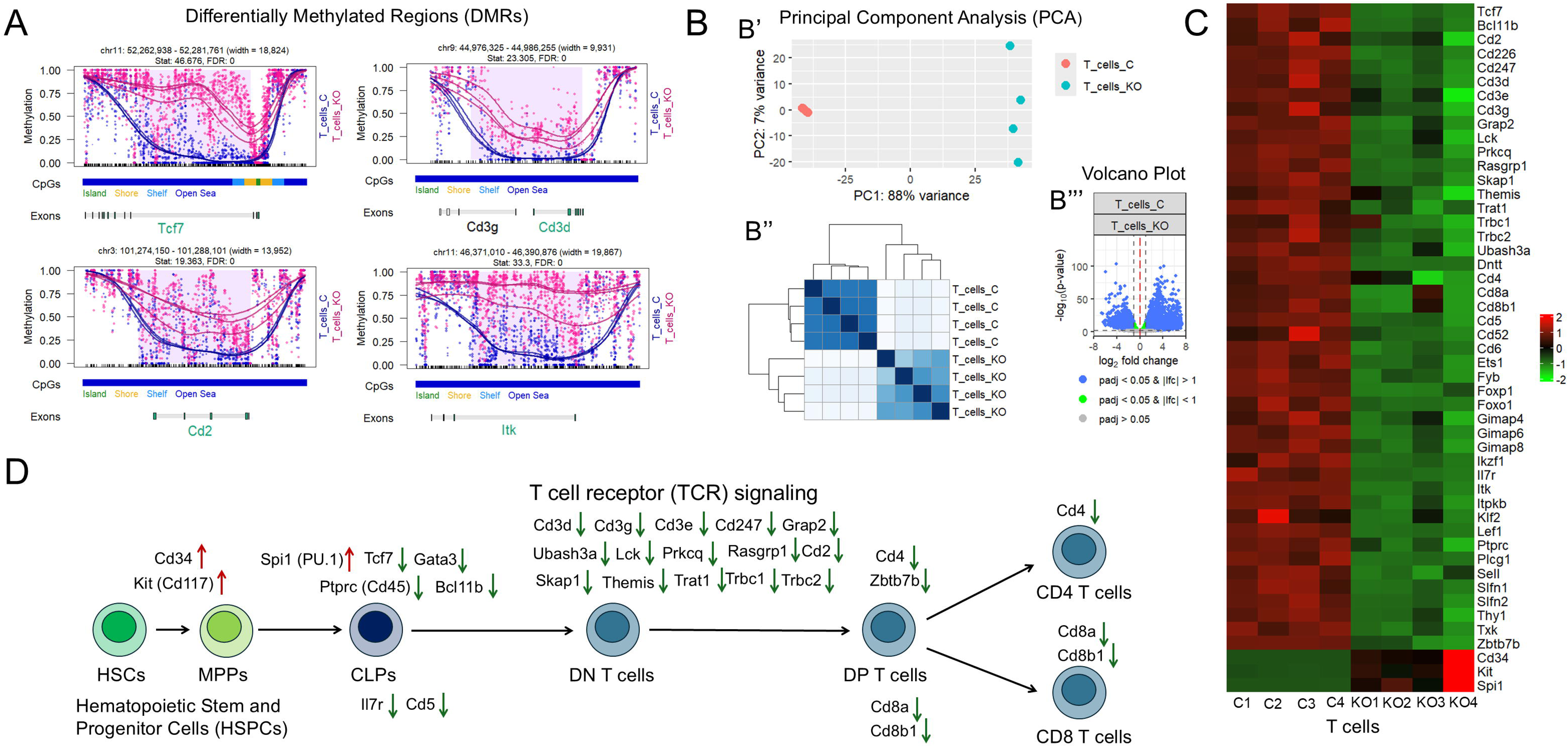
The development of T cells is controlled by the TET dependent DNA demethylation pathway. **A)** Inactivation of TET enzymes results in the maintenance of high levels of methylation of genes essential for T cell development, according to DMRseq analysis (T_cells_KO - T cells isolated from the spleen of mice treated with tamoxifen; T_cells_C - T cells isolated from the spleen of control mice). **B)** PCA (B’), heatmap (B’’), and volcano plot (B’’’) indicate a significant difference in gene expression between TET-deficient (T_cell_KO) and control (T_cell_C) T cells. **C)** The heatmap shows that maintaining high levels of T cell gene methylation leads to a decrease in their expression. **D)** Many genes whose expression is reduced in TET-deficient T cells are necessary for their specification and maturation/function. At the same time, the expression of genes that are markers of HSPCs (Cd34, Cd117) is increased in TET-deficient T cells. Taken together, these results indicate the underdevelopment of these cells. CLP – common lymphoid progenitors, DN – double negative, DP – double positive

### The TET-dependent DNA demethylation pathway controls the expression of many genes important for maturation/function, but not specification, of monocytes and granulocytes

To investigate how TET enzyme inactivation affects the development of monocytes and granulocytes at the molecular level, we isolated Cd11b positive (Cd11b^+^) cells from the bone marrow of very sick tamoxifen-treated TET deficient (KO: 11.5±2.1×10^6^ cells) mice and control (C: 11.6±1.7×10^6^) mice. Monocytes and granulocytes are short-lived, and therefore we can analyze methylome (WGBS) and transcriptome (RNA-seq) of cells that originated from TET-deficient HSPCs. Our WGBS data indicated that TSSs (±1,000 bp) of almost all monocyte and granulocyte genes we previously identified in our study are located in DMRs with higher methylation levels in Cd11b^+^ cells isolated from TET-deficient mice compared to controls (**Fig. 5A**, **Supplementary Data S2, S9**). Our RNA-seq data indicated that inactivation of TET enzymes significantly affects gene expression in isolated cells (**Fig. 5B, Supplementary Data S9**). We found that increased methylation of monocyte and granulocyte gene TSSs (±1,000 bp) in Cd11b^+^ TET-deficient cells leads to reduced expression of almost all these genes (**Supplementary Data S9**). In addition to these genes, we found reduced expression of approximately 60 other genes important for monocyte and granulocyte maturation/function (**Fig. 5C, Supplementary Data S9**). The TSSs of many of these genes are located in DMRs with increased methylation, which likely resulted in their reduced expression (**Supplementary Data S9**). We also attempted to identify the signaling cascades in which genes with increased or reduced expression are involved. Most of the genes whose expression was increased in TET-deficient monocytes/granulocytes are involved in cellular metabolism, while many genes whose expression is reduced are responsible for innate immunity, granulocyte (neutrophil) degranulation, and are necessary for cell motility (**Fig. 5D**). It should be noted that we observed increased expression of genes that promote monocyte/granulocyte specification (*Irf8*, logFC = 2.29, FDR = 3.8*10^-5^; *Klf4*, logFC = 1.58, FDR = 9.4*10^-4^; *Cebpa*, logFC = 0.89, FDR = 5.4*10^-4^). Thus, while there are no barriers to monocyte and granulocyte specification, their maturation/function may be impaired.

**Figure 5.**
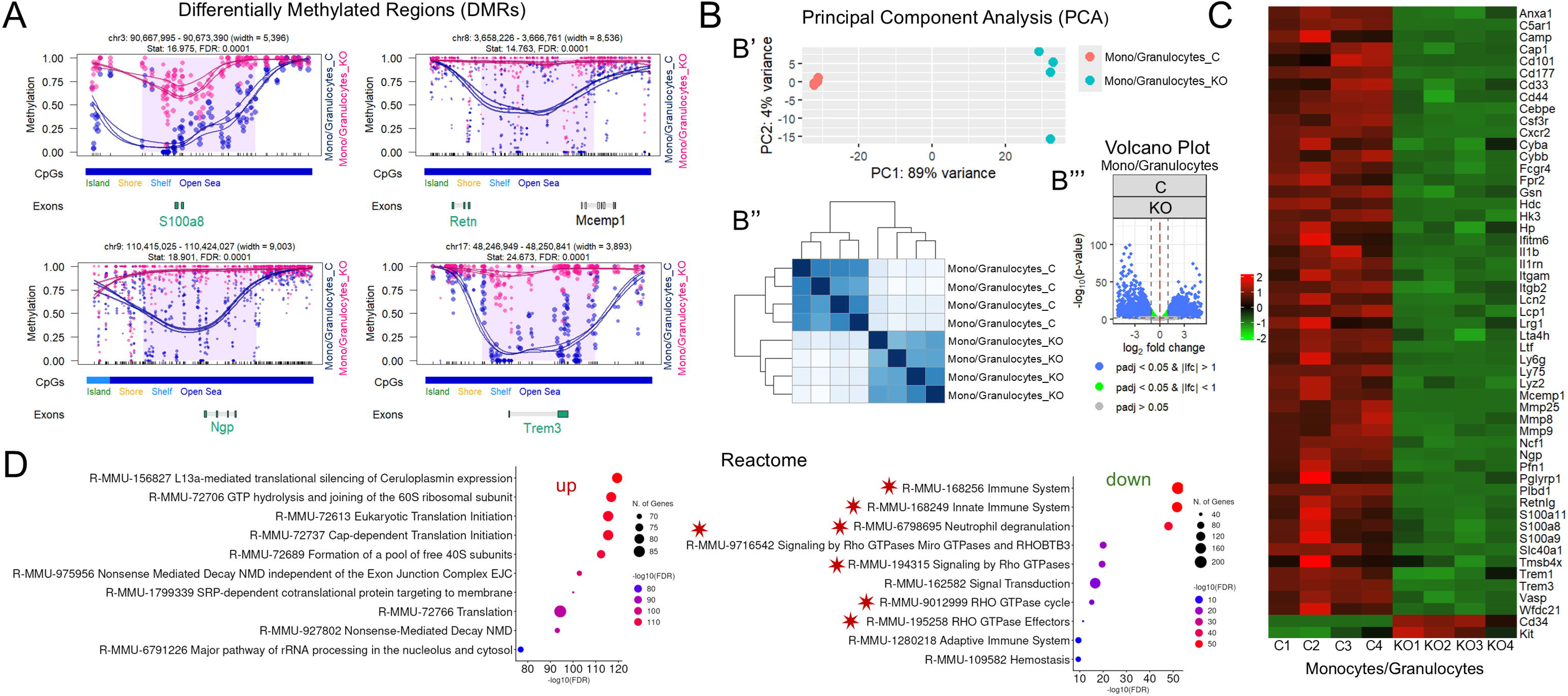
Inactivation of TET enzymes in HSPCs prevents demethylation and expression of genes essential for monocyte/granulocyte maturation/function. **A)** The panel shows examples of genes important for monocyte/granulocyte maturation/function whose methylation remained high in TET-deficient monocytes and granulocytes. **B)** Inactivation of TET enzymes significantly alters gene expression in monocytes/granulocytes, according to PCA (B’), heatmap (B’’), and volcano plot (B’’’). **C)** The heatmap shows that expression of many genes essential for monocyte and granulocyte maturation/function is reduced in TET-deficient Cd11b^+^ cells. **D)** Reactome enrichment analysis of genes with increased (up) and decreased (down) expression (Counts Per Million [CPM] >100) by a factor of 1.5 was performed to identify important signaling cascades.

### TET-deficient HSPCs fail to differentiate and eventually accumulate due to the lack of demethylation and activation of genes necessary for the specification and maturation of various blood cell types during hematopoiesis

Cd117 (Kit) is expressed on HSPCs and early myeloid/lymphoid progenitors and is thus an ideal tool for studying the early stage of hematopoiesis. Our findings indicated that inactivation of the TET-dependent DNA demethylation pathway leads to a decrease in the number of mature blood cells and an increase in the number of HSPCs (**Fig. 3**). To investigate this phenomenon, we isolated Cd117 positive (Cd117^+^) cells from the bone marrow of tamoxifen-treated (Cd117_KO) and control (Cd117_C) mice. In agreement with our flow cytometry data, the number of Cd117^+^ cells isolated from the bone marrow of TET-deficient mice was almost twice the number of those isolated from controls (Cd117_KO: 15.6±1.9×10^6^ vs. Cd117_C: 7.8±1.1×10^6^; P-value=0.0024). Isolated Cd117^+^ cells were used in WGBS and RNA-seq. We found many DMRs with increased methylation levels in KO vs. C Cd117^+^ cells containing TSSs (±1,000 bp) of the genes we were studying (**Fig. 6A, Supplementary Data S10**). It should be noted that the level of methylation in DMRs is high for both experimental and control Cd117^+^ cells (**Fig. 6A, Supplementary Data S10**). This is explained by the fact that the Cd117 cell pool we isolated included cells at different stages of differentiation: from HSPCs, in which these genes are hypermethylated and inactive, to myeloid/lymphoid progenitors, in which these genes are likely hypomethylated and active. The higher level of methylation in TET-deficient Cd117^+^ cells indicates that they lack DNA demethylation. The absence of TET enzyme activity in TET-deficient Cd117^+^ cells also significantly affected gene expression (**Fig. 6B, Supplementary Data S10**). The expression of many genes involved in hematopoiesis was reduced in TET-deficient Cd117^+^ cells (**Fig. 6C, Supplementary Data S10**). TSSs (±1,000 bp) of many of these genes are located in DMRs with increased methylation that likely suppressed their expression (**Supplementary Data S10**). By analyzing gene expression depending on the cell type, we noticed that genes required for HSPC, CLP, and CMP development do not change their expression or even increase it when TET enzymes are inactivated (**Fig. 6D**, **Supplementary Data S10**). In contrast, the expression of genes necessary for the specification and maturation of erythroblasts, megakaryocytes, and B cells was significantly reduced in TET-deficient Cd117^+^ cells, which may have hindered their generation and lead to the low numbers we detected in bone marrow (**Fig. 3, 6D, Supplementary Data S10**). Interestingly, in TET-deficient Cd117^+^ cells, we observed a decrease in the expression of genes necessary for maturation, but not specification, of monocytes and granulocytes, which may not prevent the appearance of these cells, although their function could be impaired (**Fig. 6D, Supplementary Data S10**). This result is consistent with earlier observations (**Fig. 3, 5**). Analysis of Cd117^+^ cells proliferation based on EdU incorporation into DNA revealed that the Cd117_KO cells divide at half the rate of control cells (**Fig. 6E**). A reduced rate of Cd117_KO cell proliferation may be explained by the absence of erythroid precursors (erythroblasts), which have the highest proliferation rate in the bone marrow^17^. At the same time, the Cd117_KO cells exhibited remarkable viability (**Fig. 6F**). These observations are also supported by our RNA-seq data (**Supplementary Data S10**). Thus, the inability to differentiate and high viability lead to the accumulation of TET-deficient HSPCs even with the reduced proliferation rate.

**Figure 6.**
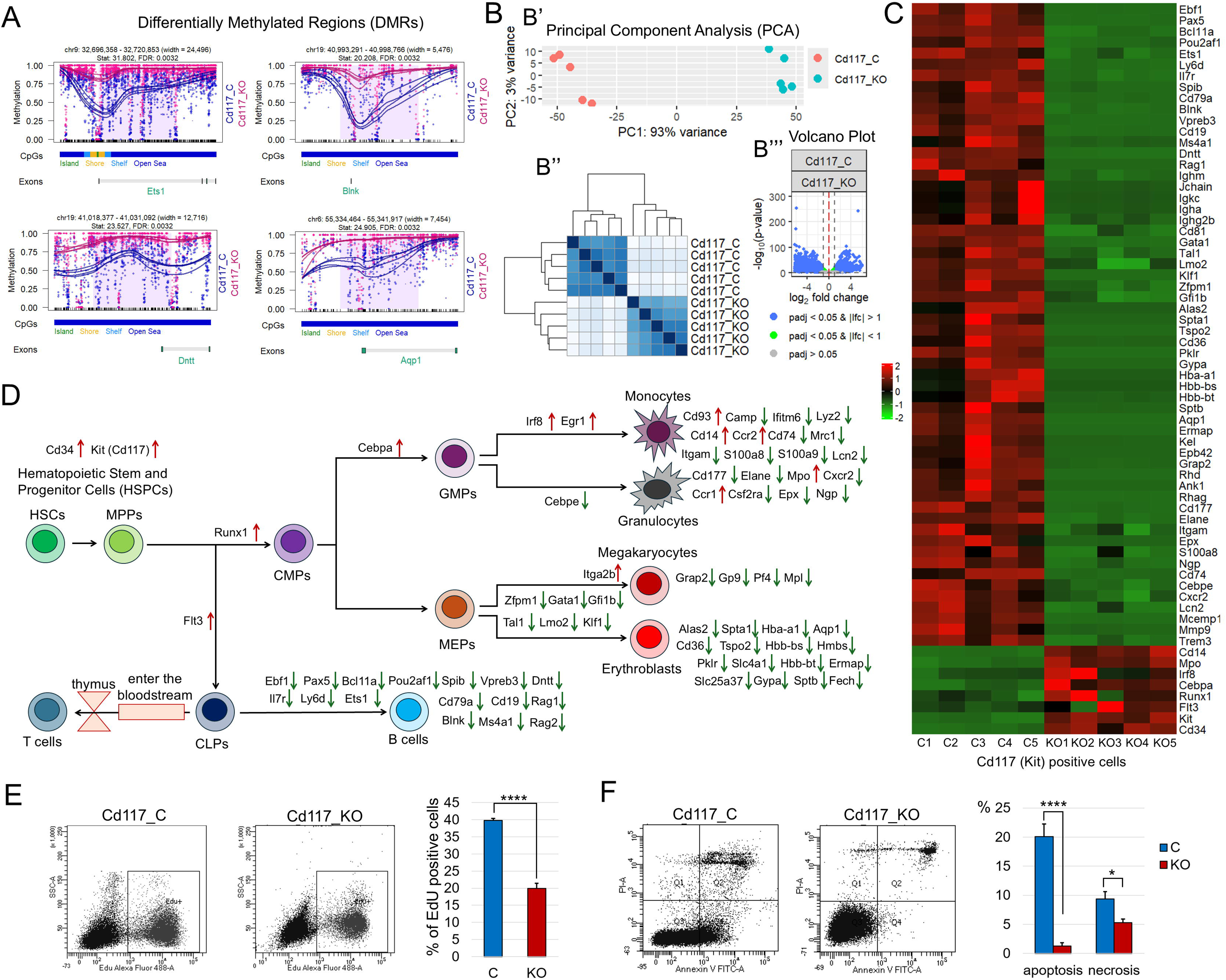
Demethylation and activation of many hematopoietic genes are suppressed early in hematopoiesis in TET-deficient experimental mice. **A)** The TET dependent DNA demethylation pathway initiates the demethylation of hematopoietic genes as early as the myeloid/lymphoid progenitor stage. DMRs were identified by comparing the methylomes of experimental (Cd117_KO) and control (Cd117_C) Cd117^+^ cells using the DMRseq Bioconductor package. **B)** Changes in methylation patterns caused by inactivation of TET enzymes lead to significant changes in gene expression in Cd117^+^ cells as evidenced by PCA (B’), heatmap (B’’), and volcano plot (B’’’). **C)** The heatmap reflects changes in the expression of many genes involved in hematopoiesis in Cd117_KO vs Cd117_C cells. **D)** The panel shows the stages of hematopoiesis and changes in the expression of genes corresponding to these stages. A green downward arrow indicates decreased gene expression in TET-deficient Cd117^+^ cells, while a red upward arrow indicates increased gene expression. **E)** To examine the rate of Cd117^+^ cell proliferation, EdU was administered intraperitonially into experimental and control mice. The bone marrow of these animals was collected after 30 minutes and used for Cd117^+^ cell isolation. Cd117^+^ cells were labeled, and the percentage (%) of cells incorporating EdU was determined using flow cytometry (n=5, ****P-value < 0.0001). **F)** The percentage (%) of apoptotic (Annexin V+/PI-; Q4) and necrotic (PI+; Q1 and Q2) Cd117^+^ cells was determined using flow cytometry (n=5, ****P-value < 0.0001, *P-value < 0.05).

## Discussion

Hematopoiesis is a complex process of generating various types of blood cells from HSPCs that is tightly regulated by multiple signaling cascades^1–3^. In this study, we demonstrated that a unique feature of these signaling cascades is that many of the genes comprising them are highly methylated in HSPCs and must undergo demethylation to become activated in mature blood cells. We further demonstrated that the TET-dependent DNA demethylation pathway is responsible for the demethylation of all these genes during HSPC differentiation, thereby exerting global control over hematopoiesis. Disrupted activity of this pathway leads to severe impairment of hematopoiesis and results in hematological malignancies. Our findings indicate that these malignancies are not caused by uncontrolled proliferation of TET-deficient HSPCs, but rather by the inability of the HSPCs to differentiate due to a failure to demethylate the genes involved in hematopoiesis.

We discovered that many genes critical for the specification and maturation of erythroblasts, T cells, and B cells are highly methylated in human and mouse HSPCs. We also discovered that many genes required for the maturation, but not for the specification, of monocytes and granulocytes are highly methylated in the HSPCs. We demonstrated that TET enzymes demethylate all these genes, a process essential for their activation in mature blood cell types. All these genes undergo demethylation during hematopoiesis in accordance with a “specific rule”: during the differentiation of HSPCs into a specific blood cell type, only the genes required for that cell type are selectively demethylated by TET enzymes, while genes required for other cell types remain hypermethylated. The exception to this rule is found in erythroblasts, during whose differentiation virtually the entire genome undergoes demethylation. It should be noted that global DNA demethylation during mouse erythropoiesis has been observed previously using reduced representation bisulfite sequencing (RRBS)^18^. Since the development of erythroblasts, T cells, and B cells requires the demethylation of numerous genes critical for their specification and maturation, the inactivation of TET enzymes results in reduced cell counts for these lineages in mice. This effect is particularly pronounced in the erythrocyte/erythroblast population, given the short lifespan of these cells. The absence of TET enzymes does not, however, impede the generation of monocytes and granulocytes. Nevertheless, the function of these blood cells may be impaired, as TET-deficient monocytes and granulocytes exhibit reduced expression of many genes governing their inflammatory response, motility, and degranulation. It should be also noted that the interaction of various types of blood cells is required for hematopoiesis^1–3^. Therefore, disruptions in the development of one blood cell type can affect the development of others. We found that there was more than one blood cell type with impaired development, which should undoubtedly affect hematopoiesis.

Our findings indicate that TET enzymes selectively demethylate genes essential for the development of a specific blood cell type during hematopoiesis. How can this selectivity be explained? Another paradox lies in the fact that TET enzymes, which are responsible for DNA demethylation, are unable to bind to hypermethylated DNA^9,19–24^. TET enzymes resolve both issues by forming a complex with specific transcription factors, which, in turn, bind to the transcription start sites (TSSs) of target genes to facilitate their demethylation and activation^25–35^. At least three such transcription factors (Spi1 [PU.1], Wt1, and Runx1) that can bind to TET (Tet2) enzymes and deliver them to TSSs of genes responsible for hematopoiesis have already been discovered^31,32,36^. Thus, during the differentiation of HSPCs into a specific blood cell type, a unique set of transcription factors must exist to govern the precise demethylation of genes required exclusively for that particular cell type. One such set is responsible for the development of erythrocytes/erythroblasts, another for the development of T cells, and so on. Collectively, they constitute the TET-dependent DNA demethylation pathway that exerts global control over hematopoiesis. The identification of these transcription factors is critical to understanding the molecular mechanisms by which the TET-dependent DNA demethylation pathway exerts global control over hematopoiesis. It should be noted that the TET-dependent DNA demethylation pathway plays a similar and important role in the development of other tissues including the retina and bones^37–39^. Thus, similar TET-dependent epigenetic mechanisms are likely responsible for the development of many tissues. Energetically, we posit it is likely more advantageous to demethylate the necessary genes during stem cell differentiation rather than to specifically methylate a multitude of unnecessary genes (**Fig. 7A**).

**Figure 7.**
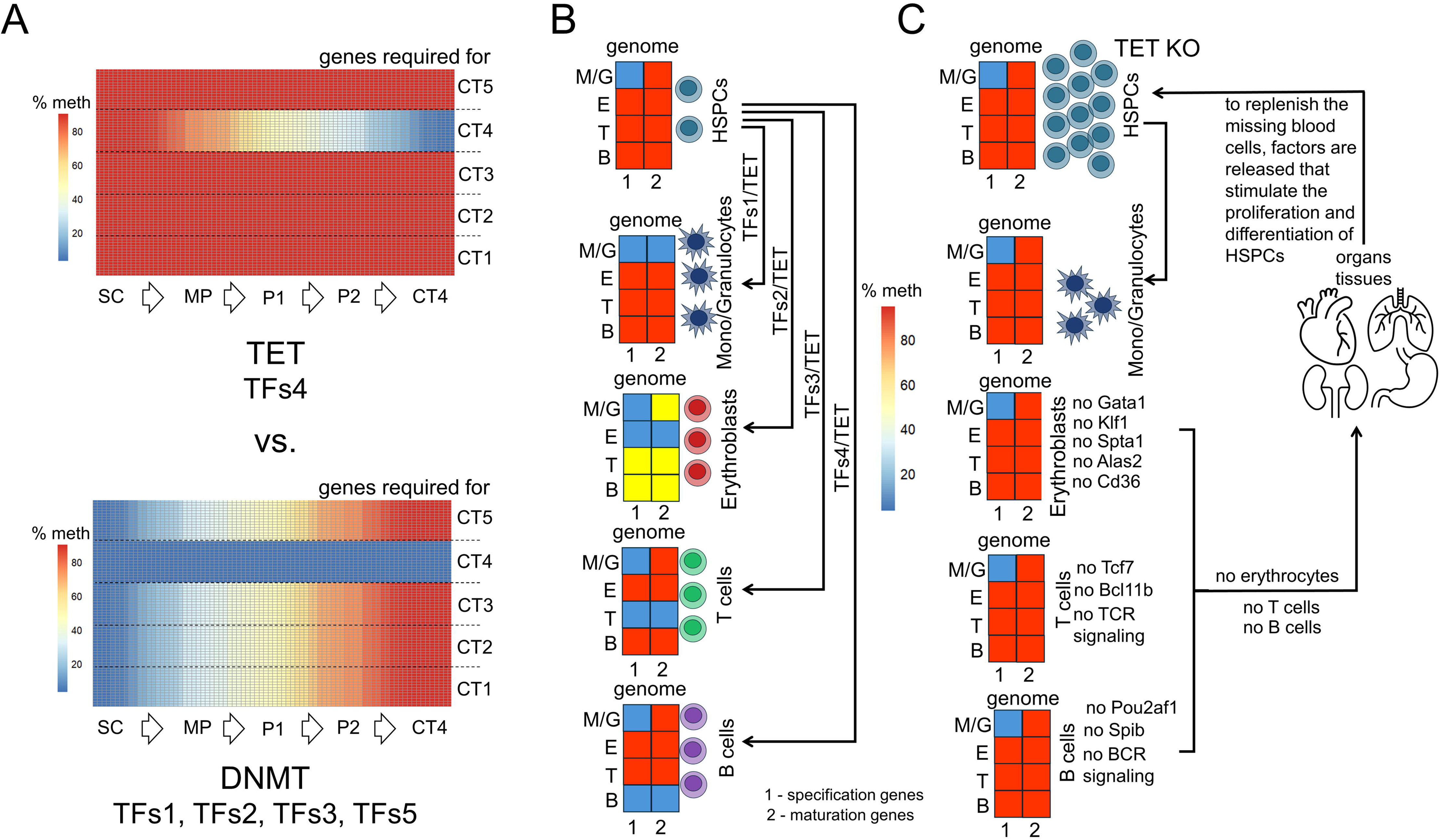
The process of TET-dependent DNA demethylation has advantages over DNMT-dependent DNA methylation in tissue development. **A)** If many genes of various cell types are hypermethylated in stem cells (SC), then the differentiation of stem cells into a specific cell type (on the panel it is marked as CT4) requires a single group of transcription factors (TFs4) to demethylate and activate those genes in a TET-dependent manner. However, if the genes necessary for the development of various cell types exhibit low levels of methylation in stem cells, then DNMT enzymes will likely require the assistance of several groups of transcription factors to specifically methylate the unnecessary genes during stem cell differentiation to ensure their inactivity in a specific cell type (CT4). This option may be energetically unfavorable for a cell. **B)** For TET-dependent demethylation of hematopoietic genes during the differentiation of HSPCs into various types of blood cells, many groups of transcription factors (TFs) are needed that determine the specificity of TET enzyme’s activity. **C)** Inactivation of TET enzymes leaves genes important for the specification and maturation/function of various blood cell types highly methylated and inactive, preventing TET-deficient HSPCs from differentiating into these cell types. The absence of blood cells necessary for normal bodily function stimulates the release of factors that promote TET-deficient HSPC proliferation but not differentiation, triggering a positive feedback loop leading to severe pathology and death of the organism.

Disrupted activity of the TET-dependent DNA demethylation pathway, caused by mutations in its genes, leads to the development of various types of leukemia^8,9^. The genes encoding TET enzymes are the ones most frequently mutated in these hematological malignancies^8,9^. The results of this study, along with numerous publications, indicate that the inactivation of TET genes in mice leads to the development of the same symptoms observed in patients suffering from leukemia^40–46^. However, it was previously hypothesized that TET enzymes suppress the development of leukemia through mechanisms entirely unrelated to DNA demethylation^40,46^. Our findings, for the first time, provide an explanation of how the primary function of TET enzymes is essential for hematopoiesis and how its disruption can lead to leukemia. We discovered that lack of TET-dependent demethylation of key hematopoietic genes prevents HSPC differentiation. The inability of the TET-deficient HSPCs to differentiate results in the depletion of the pool of blood cells critical for the body’s vital functions (etc., erythrocytes/erythroblasts, T cells, B cells), which would be expected to trigger the secretion of factors that promote the proliferation of HSPCs with their subsequent differentiation into missing cells (**Fig. 7B, 7C**). We found that the TET-deficient HSPCs proliferate and are highly viable, yet—as noted above—they are unable to differentiate. This results in HSPC accumulation, while the pool of essential blood cells remains unreplenished. All these processes trigger a positive feedback loop that can lead to lethality within a short timeframe (**Fig. 7C**). Since the proper functioning of the TET-dependent DNA demethylation pathway requires the activity of numerous transcription factors responsible for the specificity of gene demethylation during hematopoiesis, mutations in these transcription factors would be expected to give rise to a range of blood disorders, including leukemia. The severity of these diseases may depend on the number of important hematopoietic genes for the specific TET-dependent demethylation of which these transcription factors are responsible.

In conclusion, we have demonstrated for the first time that the development of virtually all types of blood cells depends on the activity of a single epigenetic pathway, the TET-dependent DNA demethylation pathway. This unique phenomenon is explained by the high level of methylation in HSPCs of an enormous number of genes responsible for the development of these blood cell types. This underlies the ability of the TET-dependent DNA demethylation pathway to exert global control over hematopoiesis. Transcription factors, which are responsible for the specificity of TET enzymes during the differentiation of HSPCs into various blood cell types, must play a crucial role in the TET-dependent DNA demethylation pathway’s activity. Studies identifying these transcription factors will be necessary for ultimately understanding how the TET-dependent DNA demethylation pathway exerts global control over hematopoiesis.

## Materials and Methods

### Animals and ethical approvals

The procedures were performed in compliance with the NIH Guide for the Care and Use of Laboratory Animals, Animal Research: Reporting of In Vivo Experiments guidelines (ARRIVE), and according to the University of Miami Institutional Animal Care and Use Committee (IACUC) approved protocol. Tet1- floxed, Tet2-floxed, and Tet3-floxed mice were acquired from La Jolla Institute for Immunology (Dr. Anjana Rao Lab), crossed to produce TET triple floxed mice, and used to generate TET global conditional knockouts. To this end, TET triple floxed mice were crossed with tamoxifen-inducible R26-CreERT2 mice (strain 008463, the Jackson Laboratory, Bar Harbor, ME, USA) to create mice homozygous for TET floxed genes and heterozygous for CreERT2. These mice have the C57BL/6J genetic background. To inactivate TET genes in all cells (KO), including HSPCs, the two-month-old mice were treated with tamoxifen (T5648, MilliporeSigma, Burlington, MA, USA) dissolved in corn oil (HY-Y1888, MedChemExpress, Monmouth Junction, NJ, USA) for 5 days (100 µL, 20 mg/mL). Mice treated only with oil were considered as controls (C). Both males and females were used to address sex as a biological variable. Erythroblasts were isolated from the bone marrow of C57BL/6J mice (strain 000664, the Jackson Laboratory, Bar Harbor, ME, USA). The mice were kept in standard housing conditions on a 12-hour light/dark cycle with unrestricted access to food and water. Animals were euthanized according to the recommendations of the Panel on Euthanasia of the AVMA.

### Tissue collection

The spleen, liver, lung, femur (thigh bone), and tibia (shin bone) were collected after the TET knockout (KO) and control (C) mice were euthanized. The weight of the spleen, liver, and lungs was determined. The size of the spleen was also determined, and it was then used for T cell isolation and in fluorescence flow cytometry (FFC). The bone marrow was collected from femur and tibia bones. To prepare the serum, freshly collected blood in tubes was left undisturbed for 30 min to allow the blood to clot completely. These tubes were spun in a refrigerated centrifuge at 2,000 g for 15 min to separate the solid clot from the liquid serum. The clear, yellowish liquid (serum) from the top was then transferred to a clean, labeled tube, leaving the clot behind. The serum was stored at -80^0^C if not analyzed immediately.

### Blood cell types used in the study and their isolation

Blood cells were isolated using magnetically labeled antibodies and columns placed in a magnetic field, in accordance with the MACS separation protocols (Miltenyi Biotec, Auburn, CA USA). Mouse erythroblasts were isolated from bone marrow using anti-Ter-119 MicroBeads and MACS separation protocol (130-049-901, Miltenyi Biotec, Auburn, CA USA). To isolate mouse monocytes and granulocytes from bone marrow, we used CD11b MicroBeads UltraPure (130-126-725, Miltenyi Biotec, Auburn, CA USA). Cd117 (cKit) positive mouse cells were isolated from bone marrow using CD117 MicroBeads according to the MACS protocol (130-091-224, Miltenyi Biotec, Auburn, CA USA). T cells were isolated from the spleen in two steps. First, Pan T Cell Isolation Kit II (130-095-130, Miltenyi Biotec, Auburn, CA USA) was used that allows for T cell isolation based on negative selection. The isolation of T cells is based on depletion of magnetically labeled CD11b, CD11c, CD19, CD45R (B220), CD49b (DX5), CD105, anti-MHC-class II, and Ter-119 positive mouse cells. Then, CD90.2 MicroBeads (130-121-278, Miltenyi Biotec, Auburn, CA USA) were used for positive selection of T cells.

### EdU Cd117 (cKit) positive cell proliferation assay in vivo

The EdU (50mg/kg in 1X PBS) solution was injected intraperitoneally into experimental and control mice. After 30 minutes (short pulse), bone marrow from these mice was collected to isolate Cd117 (cKit) positive cells. To label proliferating Cd117 (cKit) positive cells we used ClickTech EdU Cell Proliferation Kit 488 for FC IV (BCK488-IV-FC-S, MilliporeSigma, St. Louis, MO, USA) according to manufacturer’s instructions. Cd117 (cKit) positive cells were run using the LSR-Fortessa-HTS instrument (BD Biosciences, San Jose, CA). The raw data were analyzed using FlowJo software (version 10.4.1, FlowJo, Ashland, OR).

### Study of surviving, apoptotic and necrotic Cd117 (cKit) positive cells in bone marrow

We used fluorescence flow cytometry (FFC) to identify and enumerate apoptotic and necrotic Cd117 (cKit) positive cells. To this end, Cd117 (cKit) positive cells were isolated from bone marrow using CD117 MicroBeads. Apoptotic and necrotic cells were detected using the Annexin V-FITC Kit (130-092-052, Miltenyi Biotec, Auburn, CA USA) and the LSR-Fortessa-HTS instrument. Data analysis was performed with FlowJo software.

### Analysis of blood cell types in the bone marrow and spleen using fluorescence flow cytometry (FFC)

Bone marrow and spleen cell suspensions were stained using the following antibodies: anti-Gr-1 (130-112-311, Miltenyi Biotec), anti-CD11b (130-113-797), anti-CD117 (130-111-694, Miltenyi Biotec), anti-CD71 (130-120-809, Miltenyi Biotec), anti-Ter-119 (130-117-538, Miltenyi Biotec), anti-CD8 (130-111-710, Miltenyi Biotec), anti-CD4 (130-116-509, Miltenyi Biotec), anti-CD19 (130-112-035, Miltenyi Biotec), and anti-CD45R (B220) (130-110-845, Miltenyi Biotec). Cell surface antibody (1:50) staining was performed as previously described^47^. The samples were analyzed using the LSR-Fortessa-HTS instrument and FlowJo software.

### DNA and RNA isolation and quality control

The genomic DNA (gDNA) was isolated from mouse monocytes/granulocytes, erythroblasts, T cells, and Cd117 (cKit) positive cells using the DNeasy Blood and Tissue Kit (#69504, Qiagen, Hilden, Germany). Total RNA was isolated from these cells using RNeasy Plus Mini Kit (#74134, Qiagen, Hilden, Germany)^48^. The quality and quantity of gDNA and RNA were assessed using the NanoDrop One Spectrophotometer and Qubit 4 Fluorometer (ThermoFisher Scientific, Waltham, MA, USA). To assess gDNA integrity, we used the 4200 TapeStation system (Agilent Technologies, Santa Clara, CA, USA). To assess RNA integrity, we used the 2100 Bioanalyzer Instrument (Agilent Technologies, Santa Clara, CA, USA). RNA samples with an RNA Integrity Number (RIN) score of 8 or higher were used to prepare RNA□seq libraries.

### Whole genome bisulfite sequencing (WGBS) library preparation, sequencing, and data analysis

To create WGBS libraries, xGen Methyl□Seq DNA Library Prep Kit was used according to manufacturer’s instructions (#10009860, Integrated DNA Technologies, Coralville, IA, USA). The quality and quantity of the WGBS libraries were assessed using Qubit 4 Fluorometer and the 4200 TapeStation system. The WGBS libraries were sequenced on the Illumina NovaSeq X Plus using 2□×□150 paired end configuration. The generated FASTQ files have been deposited in the BioProject database (ncbi.nlm.nih.gov/bioproject/) and can be accessed using the accession number PRJNA1403508. To evaluate the quality of FASTQ data, FastQC (v0.11.8) quality control tool was used. TrimGalore (v0.4.5) with the following parameters: e 0.1, stringency 6, length 20, nextseq 20, three_prime_clip_R1 10, clip_R2 10, was used for preprocessing our FASTQ files. To align paired FASTQ reads and determine the percentage of cytosine methylation, the Bismark Bisulfite Read Mapper and Methylation Caller program and GENCODE M25 (GRCm38.p6) Mus musculus reference (mm10) genome were used^49^. We have applied the DSS Bioconductor package, to calculate the average base coverage of the mouse genome^50^. The average base coverage of the mouse genome was around 10X or more. To perform sample clustering, identify differentially methylated cytosines (DMCs), and identify genomic regions exhibiting distinct average levels of DNA methylation within the same genome we used the methylKit Bioconductor package^51^. To identify differentially methylated regions (DMRs), we used DMRseq Bioconductor package^52^. The annotatr Bioconductor package was used to annotate our data^53^. To find transcription start sites (TSS) ±1000 bp located in DMRs, we used the EPDnew database^54^. The ComplexHeatmap Bioconductor package and pheatmap R package were used to generate heatmaps.

### Bulk RNA-seq library preparation, sequencing, and data analysis

To prepare RNA□seq libraries, the mRNA-seq library preparation Kit (RK20302, ABclonal, Woburn, MA, USA) was used according to manufacturer’s instructions. These RNA□seq libraries were sequenced on the Illumina NovaSeq X Plus using 2□×□150 paired end configuration. The FASTQ files were uploaded to the BioProject database (ncbi.nlm.nih.gov/bioproject/) and can be accessed using the accession number PRJNA1403508. A standard pipeline including the STAR RNA-seq aligner, HTseq package, and GENCODE M25 (GRCm38.p6) Mus musculus reference genome (mm10) was used to calculate how many reads overlap each of the mouse genes^55,56^. The edgeR, DESeq2, and ViDGER

Bioconductor packages were used to perform the differential gene expression analysis, principal component analysis (PCA), sample clustering and generate volcano plots^57,58^. We used ShinyGO (ver. 0.85, http://bioinformatics.sdstate.edu/go/) to identify important Reactome signaling cascades and biological processes.

### Preparation of serum samples

Serum samples were thawed and concentrations were measured using a Bradford assay.10 µL of serum were taken from each sample and loaded onto a High Select Top14 Most Abundant Proteins mini depletion spin column according to manufacturer instructions (A36369, ThermoFisher Scientific). Following depletion, the concentration of protein in each sample was again evaluated. 20 µg were taken for digestion with trypsin. Following digestion, samples were acidified to 1% TFA and desalted using a Thermo HyperSep C18 column. 5 µg of the resulting elution was dried down and resuspended in 10 µL of 100 mM TEAB in preparation for tandem mass tag labeling (TMT) using the Thermo TMTpro plate. Label incorporation was analyzed by LC-MS/MS and spectral counting and ensured to be above 95%. Labeled samples were then combined, quenched with 5% hydroxylamine for 15 minutes, and vacuum centrifuged to dryness. The combined samples were then fractionated into 4 elutions of 12.5, 17.5, 22.5, and 50% acetonitrile, respectively, using the Pierce High-pH Reverse Phase spin column kit (#84868, ThermoFisher Scientific). Fractionated samples were vacuum centrifuged to dryness and resuspended in 20 µL of 2% acetonitrile, 0.1% formic acid in preparation for LC-MS/MS. 5 µL of each sample were injected for analysis.

### Liquid Chromatography-tandem Mass Spectrometry (LC-MS/MS)

A nanoflow ultra high performance liquid chromatograph (RSLC, Dionex, Sunnyvale, CA) coupled to an electrospray bench top orbitrap mass spectrometer (Orbitrap Exploris480, ThermoFisher Scientific) was used for tandem mass spectrometry peptide sequencing experiments. The sample was first loaded onto a pre-column (2 cm x 100 µm ID packed with C18 reversed-phase resin, 5µm, 100Å) and washed for 8 minutes with aqueous 2% acetonitrile and 0.1% formic acid. The trapped peptides were eluted onto the analytical column, (C18, 75 µm ID x 25 cm, 2 µm, 100Å, Dionex). The 135-minute gradient was programmed as: 95% solvent A (2% acetonitrile + 0.1% formic acid) for 3 minutes, solvent B (90% acetonitrile + 0.1% formic acid) from 5% to 7% in 2 minutes, 7% to 25% in 90 minutes, 25% to 60% in 20 minutes, then solvent B from 60% to 95% in 1 minutes and held at 95% for 3 minutes, followed by solvent B from 90% to 5% in 1 minute and re-equilibrate for 15 minutes. The flow rate on analytical column was 300 nl/min. Cycle time was set at 3 sec for data dependent acquisition. Spray voltage was 2000v and capillary temperature was 300 °C. The resolution for MS and MS/MS scans were set at 120,000 and 45,000 respectively. Dynamic exclusion was 15 seconds for previously sampled peptide peaks.

### LC-MS/MS data analysis

MaxQuant version 2.6.7.0) was used to identify peptides and quantify the TMT reporter ion intensities. Protein database was downloaded from Uniprot in January 2025. Up to 2 missed trypsin cleavage was allowed. Carbamidomethyl cystine was set as fixed modification and Methionine oxidation was set as variable modification. Both PSM and Prtoein FDR were set at 0.01. Match between runs feature was activated.

### Statistical analysis

The Student’s t□test was employed for experiments containing one variable. P□values and FDR equal to or less than 0.05 were considered statistically significant. Generation and analysis of next□generation sequencing (NGS) data were accomplished in□house according to ENCODE standards and pipelines.

Detailed information about the statistical analysis of the NGS and LC-MS/MS data can be found in the sections above.

### Data availability

The datasets collected and analyzed in the study are available in the BioProject database (accession number PRJNA1403508) and in the article/Supplementary Datafiles. We also used NCBI GEO and SRA datasets: human HSPCs (SRR17454687, SRR17454688, SRX142784), human CD14 monocytes (SRR14417033, SRR14417034, SRR14417035, SRR14417037), human erythroblasts (GSM5652274, GSM5652275, GSM5652276), human T cells (GSM5652279, GSM5652280, GSM5652281, GSM5652282, GSM5652283, GSM5652284), human B cells (GSM5652316, GSM5652317, GSM5652318, GSM5652319, GSM5652320), mouse HSPCs (SRR15810445, SRR15810444, GSM1274424, GSM1274425, GSM1274426), and mouse B cells (SRX2544840, SRX2544841, SRX2544844, SRX2544845).

## Acknowledgments

This study was supported in part by the NIH/NEI grant R01 EY035235 (D.I.), NIH/NEI Center Core grant P30 EY014801, Research to Prevent Blindness/Unrestricted Grant GR004596-1. We are sincerely grateful to Dr. Anjana Rao (La Jolla Institute for Immunology) for the TET triple-floxed mice she provided to us. The authors thank Charles K. Yaros for his expert assistance. We thank the Proteomics and Metabolomics Core Facility at the H. Lee Moffitt Cancer Center & Research Institute; an NCI designated Comprehensive Cancer Center (P30-CA076292) for providing us with high□quality data.

## Supporting information

**Supplementary Data S1.** The average percentage of methylation of promoters and first exons of mouse genes in the genomes of HSPCs, monocytes/granulocytes, erythroblasts, T cells, and B cells was obtained based on their location in regions identified using the methylKit Bioconductor package.

**Supplementary Data S2.** The eukaryotic promoter database (EPDnew) and DMRseq Bioconductor package were used to identify mouse genes whose transcription start sites (TSS; ±1,000 bp) are located in

Monocyte/Granulocyte vs. HSPC, Erythroblast vs. HSPC, T cells vs. HSPC, and B cells vs. HSPC DMRs.

**Supplementary Data S3.** Additional mouse genes that are demethylated during the differentiation of HSPCs into various types of blood cells were identified using the DMRseq Bioconductor package and EPDnew database.

**Supplementary Data S4.** The average percentage of methylation of promoters and first exons of human genes in the genomes of HSPCs, monocytes, erythroblasts, T cells, and B cells was obtained based on their location in regions identified using the methylKit Bioconductor package.

**Supplementary Data S5.** The EPDnew database and DMRseq Bioconductor package were used to identify human genes whose TSS ±1,000 bp are located in Monocyte vs. HSPC, Erythroblast vs. HSPC, T cells vs. HSPC, and B cells vs. HSPC DMRs.

**Supplementary Data S6.** Additional human genes that are demethylated during the differentiation of HSPCs into various types of blood cells were identified using the DMRseq Bioconductor package.

**Supplementary Data S7.** LC-MS/MS analysis of serum revealed a low number or impaired function of B cells and megakaryocytes in TET-deficient mice.

**Supplementary Data S8.** DMRs and results of differential gene expression analysis were obtained using the DMRseq and edgeR Bioconductor packages for T cells collected from the spleen of experimental and control mice.

**Supplementary Data S9.** DMRs and results of differential gene expression analysis were obtained using the DMRseq and edgeR Bioconductor packages for monocytes/granulocytes collected from the bone marrow of experimental and control mice.

**Supplementary Data S10.** DMRs and results of differential gene expression analysis were obtained using the DMRseq and edgeR Bioconductor packages for Cd117 positive cells collected from the bone marrow of experimental and control mice.

## Notes

### Competing Interest Statement

The authors have declared no competing interest.

