## Supplementary Data S3 for "The TET-dependent DNA demethylation pathway is the driving force of hematopoiesis"

### additional genes

#### T cells (mm10)

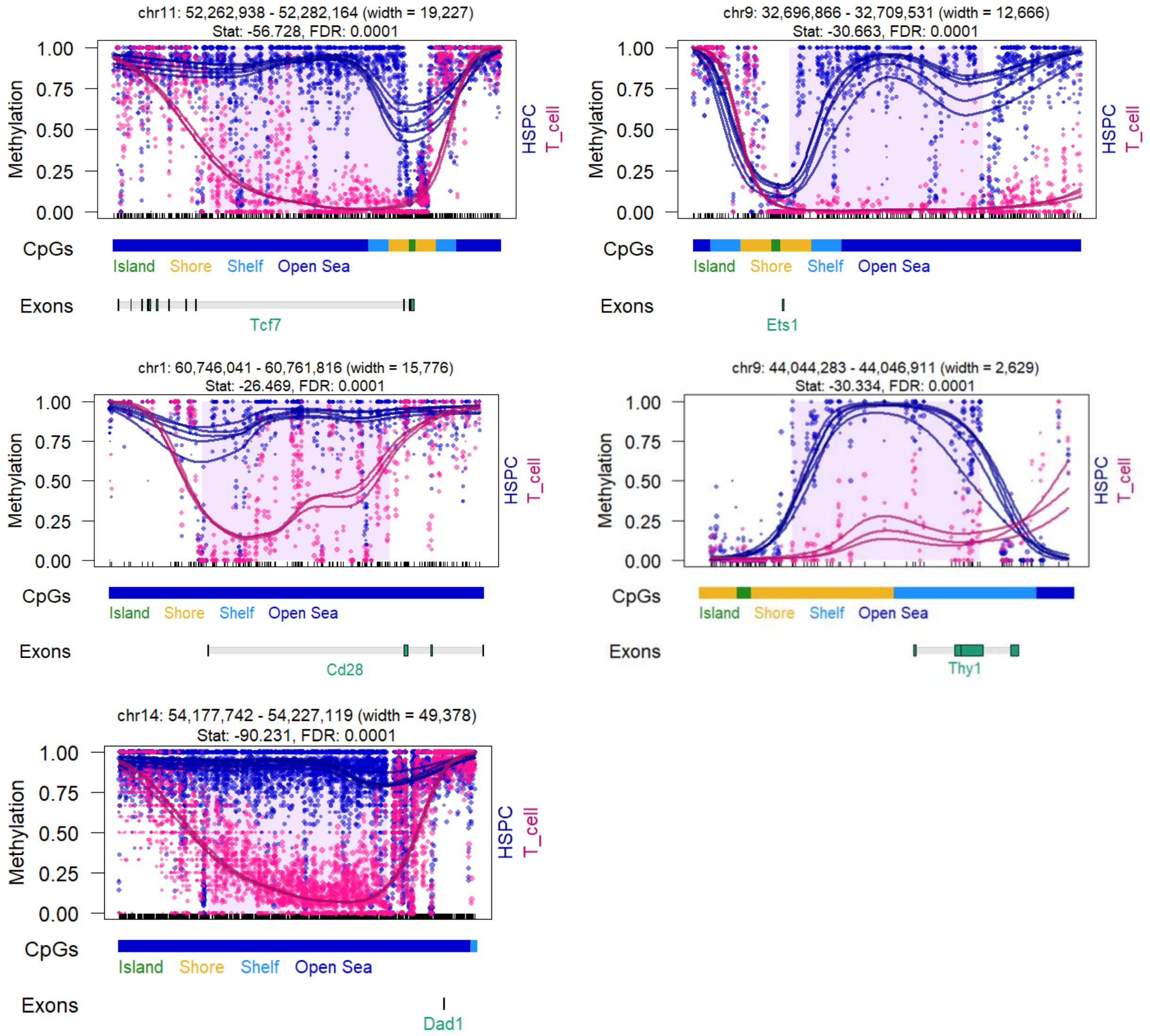


#### B cells (mm10)

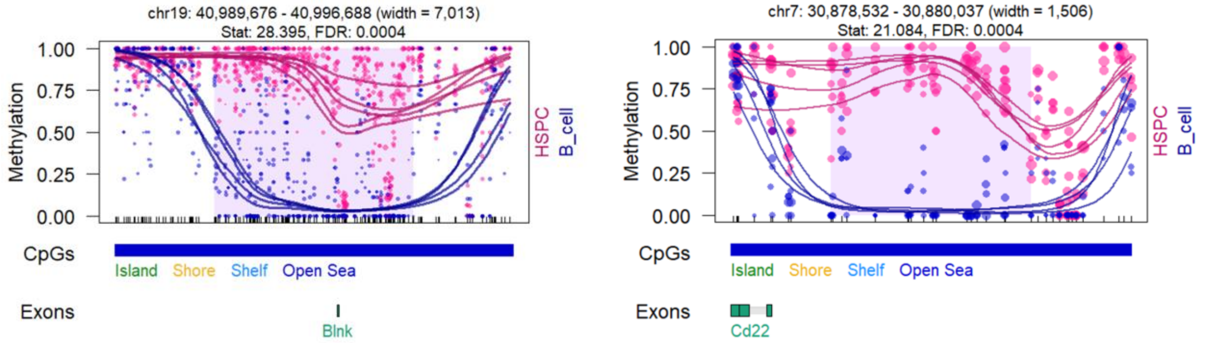

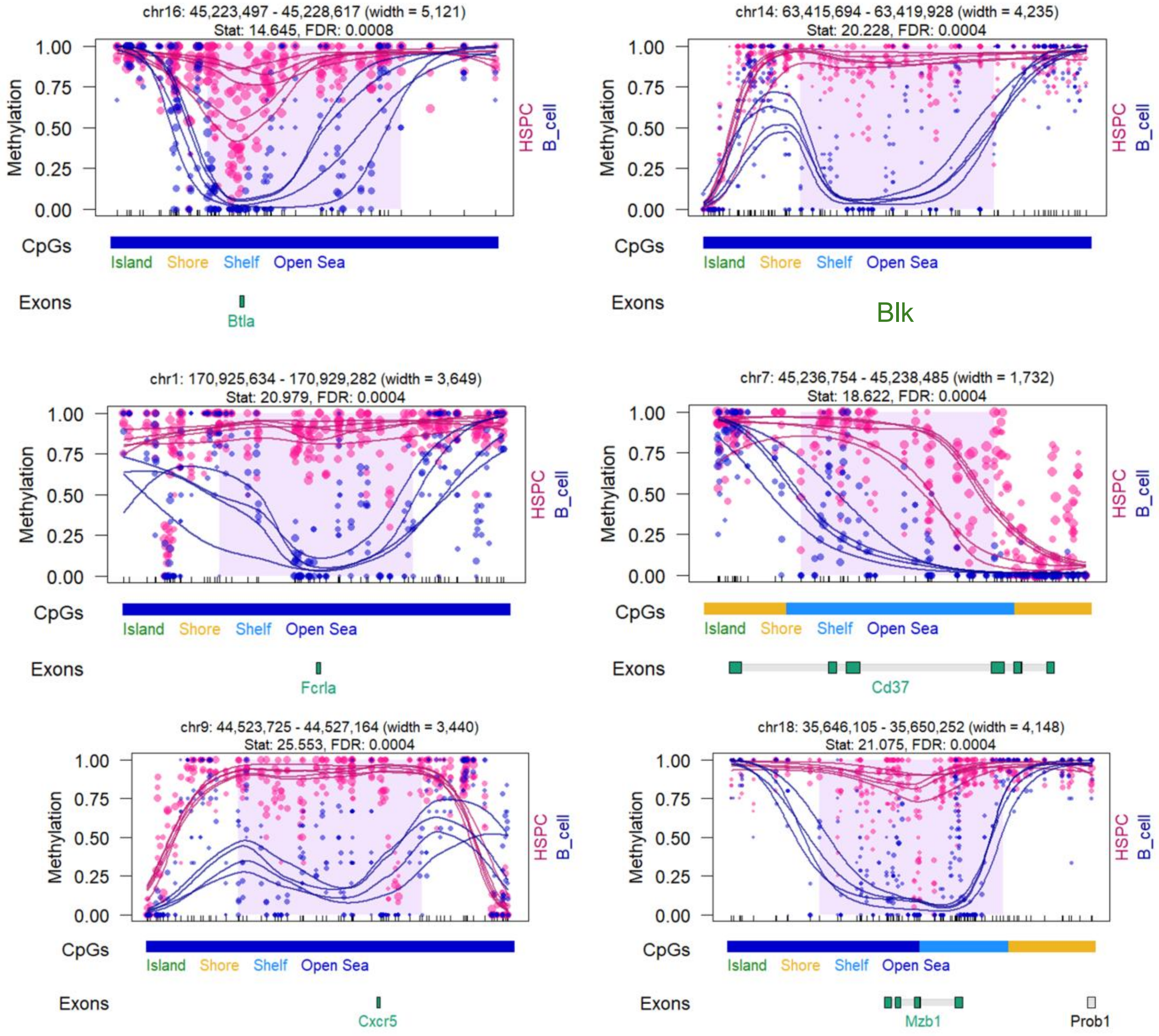
chr16: 45,223,497 - 45,228,617 (width = 5,121)
Stat: 14.645, FDR: 0.0008
Methylation
1.00
0.75
0.50
0.25
0.00
HSPC
B_cell
CpGs
Island
Shore
Shelf
Open Sea
Exons
Btla
chr14: 63,415,694 - 63,419,928 (width = 4,235)
Stat: 20.228, FDR: 0.0004
Blk
chr1: 170,925,634 - 170,929,282 (width = 3,649)
Stat: 20.979, FDR: 0.0004
Fcrla
chr7: 45,236,754 - 45,238,485 (width = 1,732)
Stat: 18.622, FDR: 0.0004
Cd37
chr9: 44,523,725 - 44,527,164 (width = 3,440)
Stat: 25.553, FDR: 0.0004
Cxcr5
chr18: 35,646,105 - 35,650,252 (width = 4,148)
Stat: 21.075, FDR: 0.0004
Mzb1
Prob1

### all genes

#### T cells (mm10)

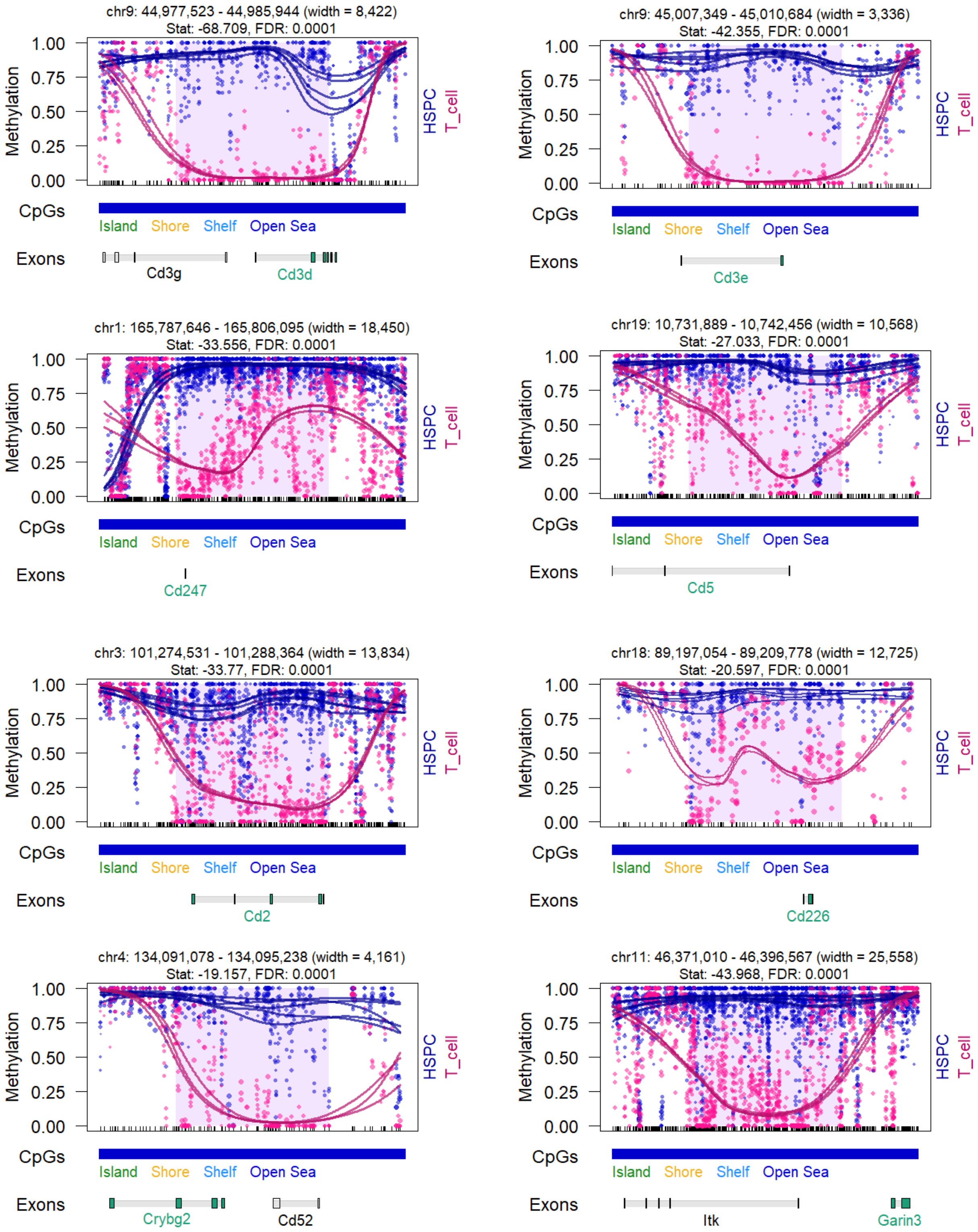

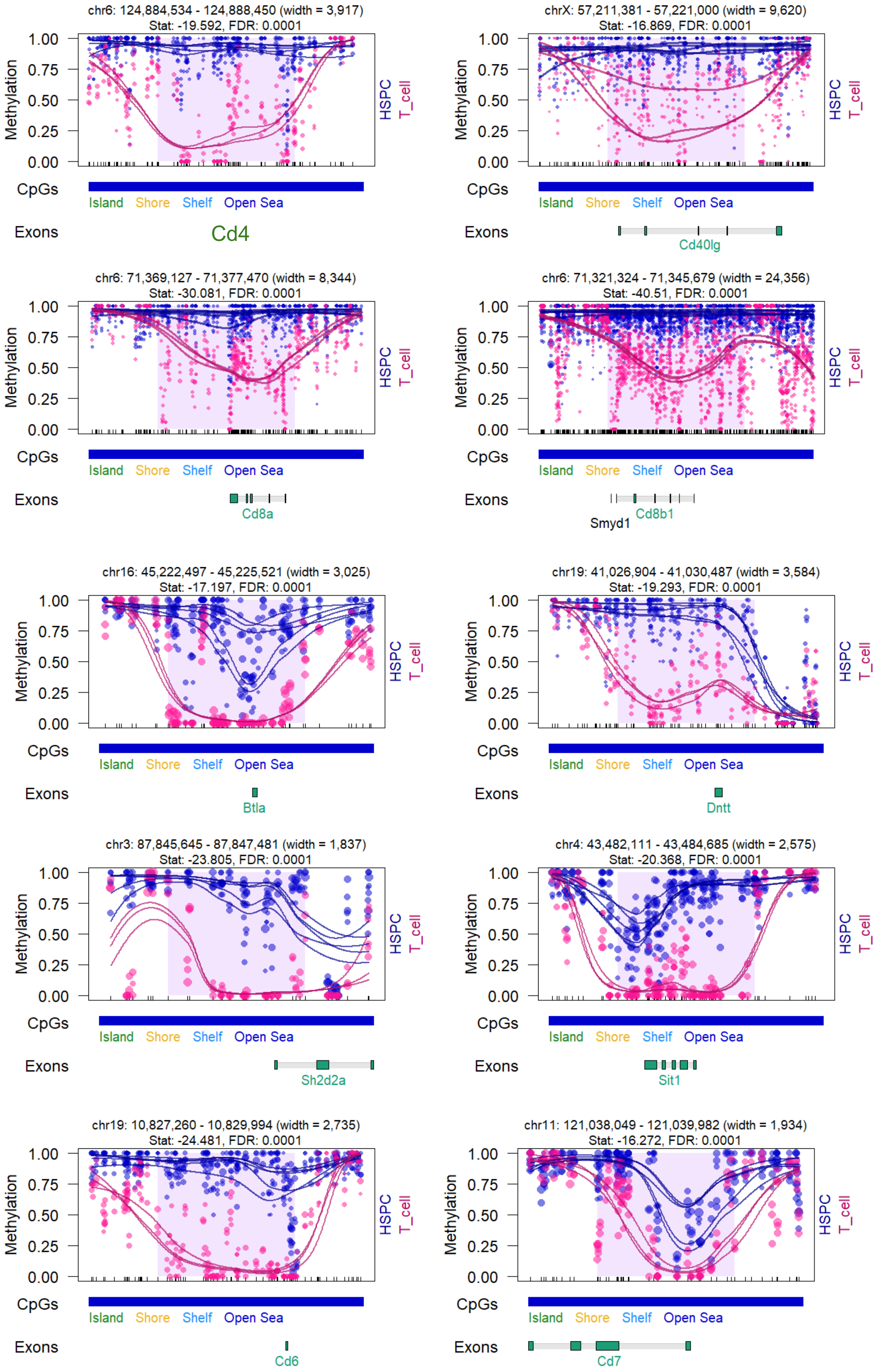
chr6: 124,884,534 - 124,888,450 (width = 3,917)
Stat: -19.592, FDR: 0.0001
Methylation
1.00
0.75
0.50
0.25
0.00
HSPC
T_cell
CpGs
Island Shore Shelf Open Sea
Exons
Cd4
chrX: 57,211,381 - 57,221,000 (width = 9,620)
Stat: -16.869, FDR: 0.0001
Methylation
HSPC
T_cell
CpGs
Island Shore Shelf Open Sea
Exons
Cd40lg
chr6: 71,369,127 - 71,377,470 (width = 8,344)
Stat: -30.081, FDR: 0.0001
Methylation
HSPC
T_cell
CpGs
Island Shore Shelf Open Sea
Exons
Cd8a
chr6: 71,321,324 - 71,345,679 (width = 24,356)
Stat: -40.51, FDR: 0.0001
Methylation
HSPC
T_cell
CpGs
Island Shore Shelf Open Sea
Exons
Smyd1
Cd8b1
chr16: 45,222,497 - 45,225,521 (width = 3,025)
Stat: -17.197, FDR: 0.0001
Methylation
HSPC
T_cell
CpGs
Island Shore Shelf Open Sea
Exons
Btla
chr19: 41,026,904 - 41,030,487 (width = 3,584)
Stat: -19.293, FDR: 0.0001
Methylation
HSPC
T_cell
CpGs
Island Shore Shelf Open Sea
Exons
Dntt
chr3: 87,845,645 - 87,847,481 (width = 1,837)
Stat: -23.805, FDR: 0.0001
Methylation
HSPC
T_cell
CpGs
Island Shore Shelf Open Sea
Exons
Sh2d2a
chr4: 43,482,111 - 43,484,685 (width = 2,575)
Stat: -20.368, FDR: 0.0001
Methylation
HSPC
T_cell
CpGs
Island Shore Shelf Open Sea
Exons
Sit1
chr19: 10,827,260 - 10,829,994 (width = 2,735)
Stat: -24.481, FDR: 0.0001
Methylation
HSPC
T_cell
CpGs
Island Shore Shelf Open Sea
Exons
Cd6
chr11: 121,038,049 - 121,039,982 (width = 1,934)
Stat: -16.272, FDR: 0.0001
Methylation
HSPC
T_cell
CpGs
Island Shore Shelf Open Sea
Exons
Cd7

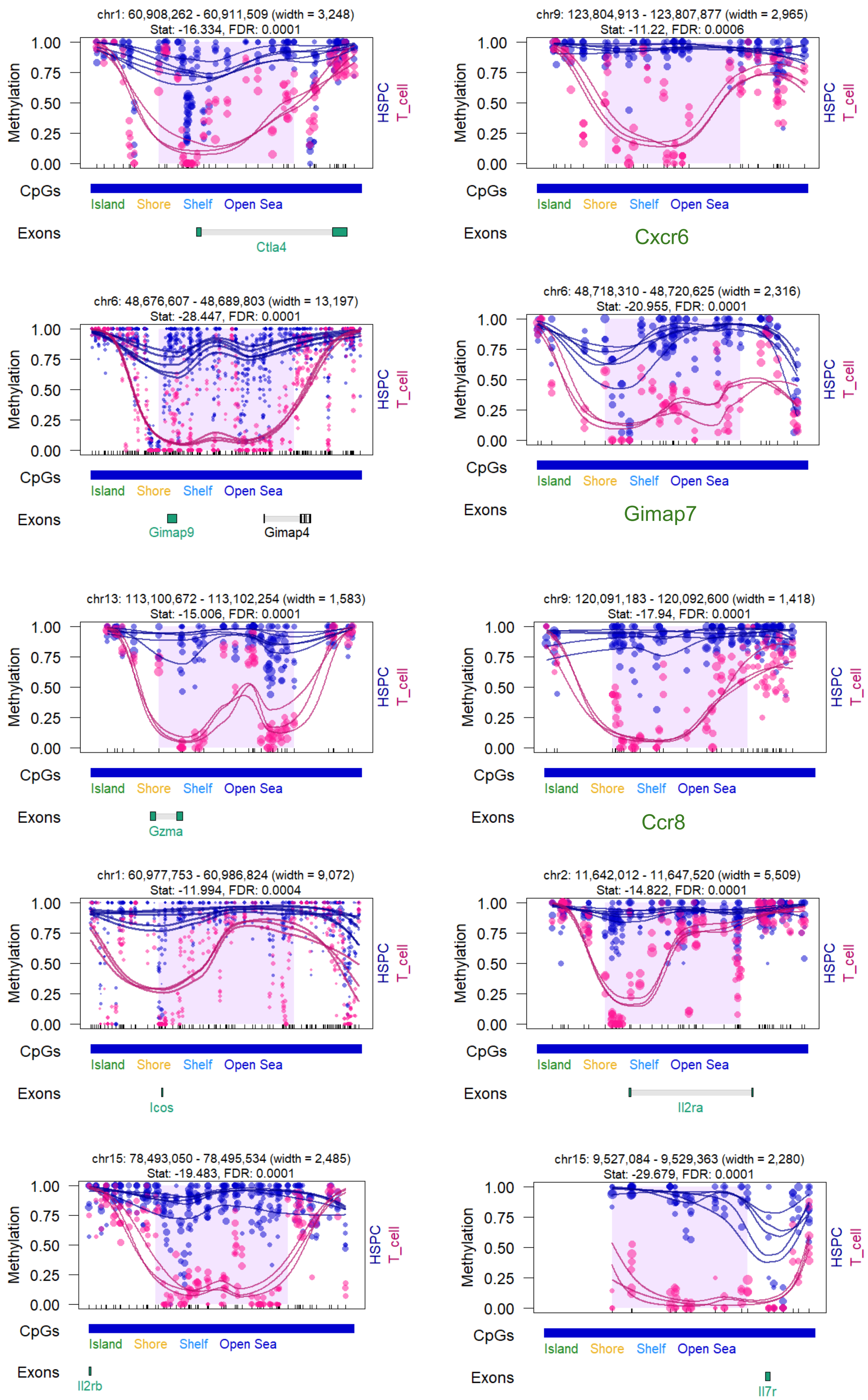
chr1: 60,908,262 - 60,911,509 (width = 3,248)
Stat: -16.334, FDR: 0.0001
Methylation
1.00
0.75
0.50
0.25
0.00
HSPC
T_cell
CpGs
Island Shore Shelf Open Sea
Exons
Ctla4
chr9: 123,804,913 - 123,807,877 (width = 2,965)
Stat: -11.22, FDR: 0.0006
Cxcr6
chr6: 48,676,607 - 48,689,803 (width = 13,197)
Stat: -28.447, FDR: 0.0001
Gimap9
Gimap4
chr6: 48,718,310 - 48,720,625 (width = 2,316)
Stat: -20.955, FDR: 0.0001
Gimap7
chr13: 113,100,672 - 113,102,254 (width = 1,583)
Stat: -15.006, FDR: 0.0001
Gzma
chr9: 120,091,183 - 120,092,600 (width = 1,418)
Stat: -17.94, FDR: 0.0001
Ccr8
chr1: 60,977,753 - 60,986,824 (width = 9,072)
Stat: -11.994, FDR: 0.0004
Icos
chr2: 11,642,012 - 11,647,520 (width = 5,509)
Stat: -14.822, FDR: 0.0001
Il2ra
chr15: 78,493,050 - 78,495,534 (width = 2,485)
Stat: -19.483, FDR: 0.0001
Il2rb
chr15: 9,527,084 - 9,529,363 (width = 2,280)
Stat: -29.679, FDR: 0.0001
Il7r

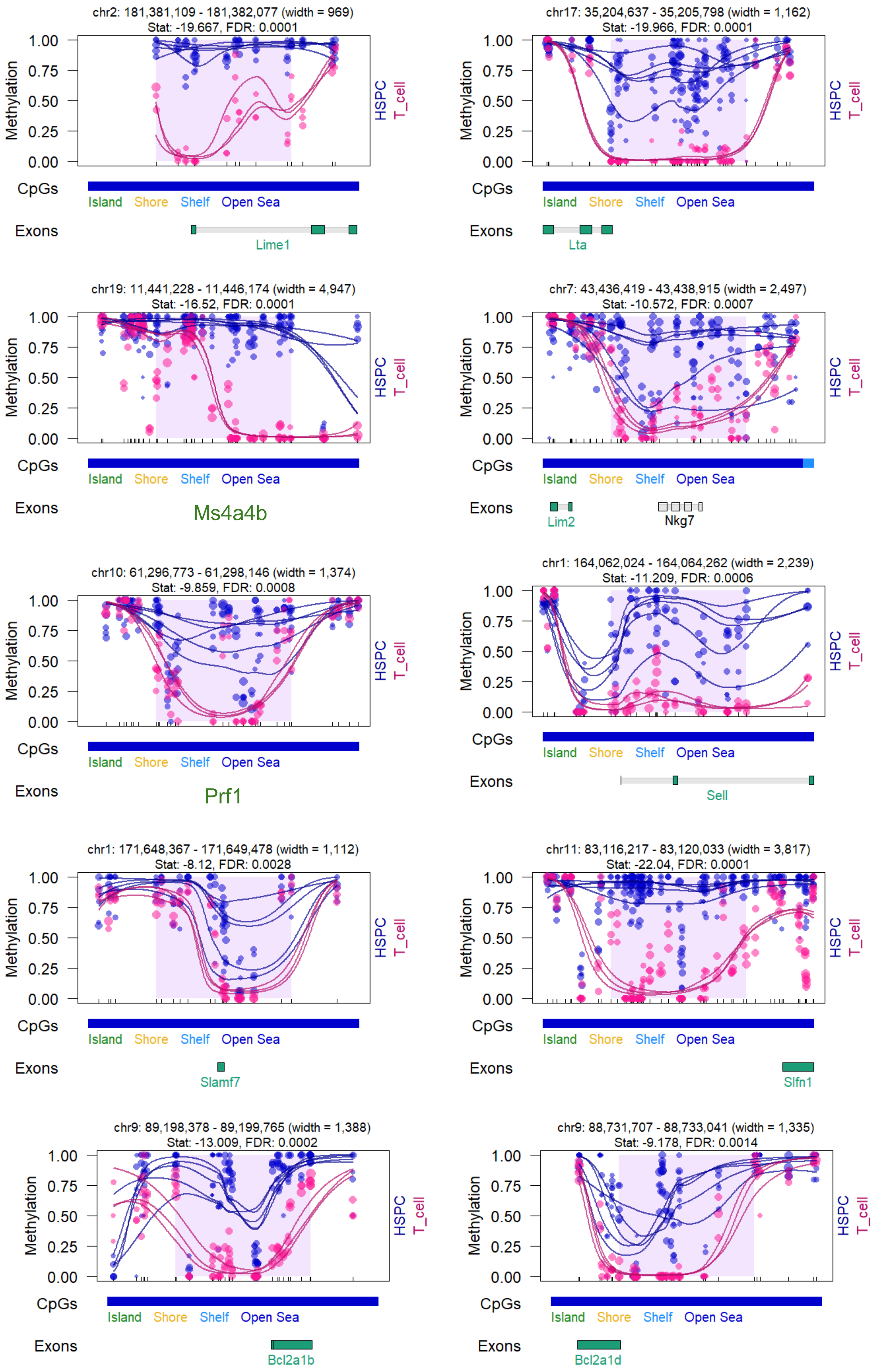

chr2: 181,381,109 - 181,382,077 (width = 969)
Stat: -19.667, FDR: 0.0001
Methylation
1.00
0.75
0.50
0.25
0.00
HSPC
T_cell
CpGs
Island
Shore
Shelf
Open Sea
Exons
Lime1
chr17: 35,204,637 - 35,205,798 (width = 1,162)
Stat: -19.966, FDR: 0.0001
Lta
chr19: 11,441,228 - 11,446,174 (width = 4,947)
Stat: -16.52, FDR: 0.0001
Ms4a4b
chr7: 43,436,419 - 43,438,915 (width = 2,497)
Stat: -10.572, FDR: 0.0007
Lim2
Nkg7
chr10: 61,296,773 - 61,298,146 (width = 1,374)
Stat: -9.859, FDR: 0.0008
Prf1
chr1: 164,062,024 - 164,064,262 (width = 2,239)
Stat: -11.209, FDR: 0.0006
Sell
chr1: 171,648,367 - 171,649,478 (width = 1,112)
Stat: -8.12, FDR: 0.0028
Slamf7
chr11: 83,116,217 - 83,120,033 (width = 3,817)
Stat: -22.04, FDR: 0.0001
Slfn1
chr9: 89,198,378 - 89,199,765 (width = 1,388)
Stat: -13.009, FDR: 0.0002
Bcl2a1b
chr9: 88,731,707 - 88,733,041 (width = 1,335)
Stat: -9.178, FDR: 0.0014
Bcl2a1d

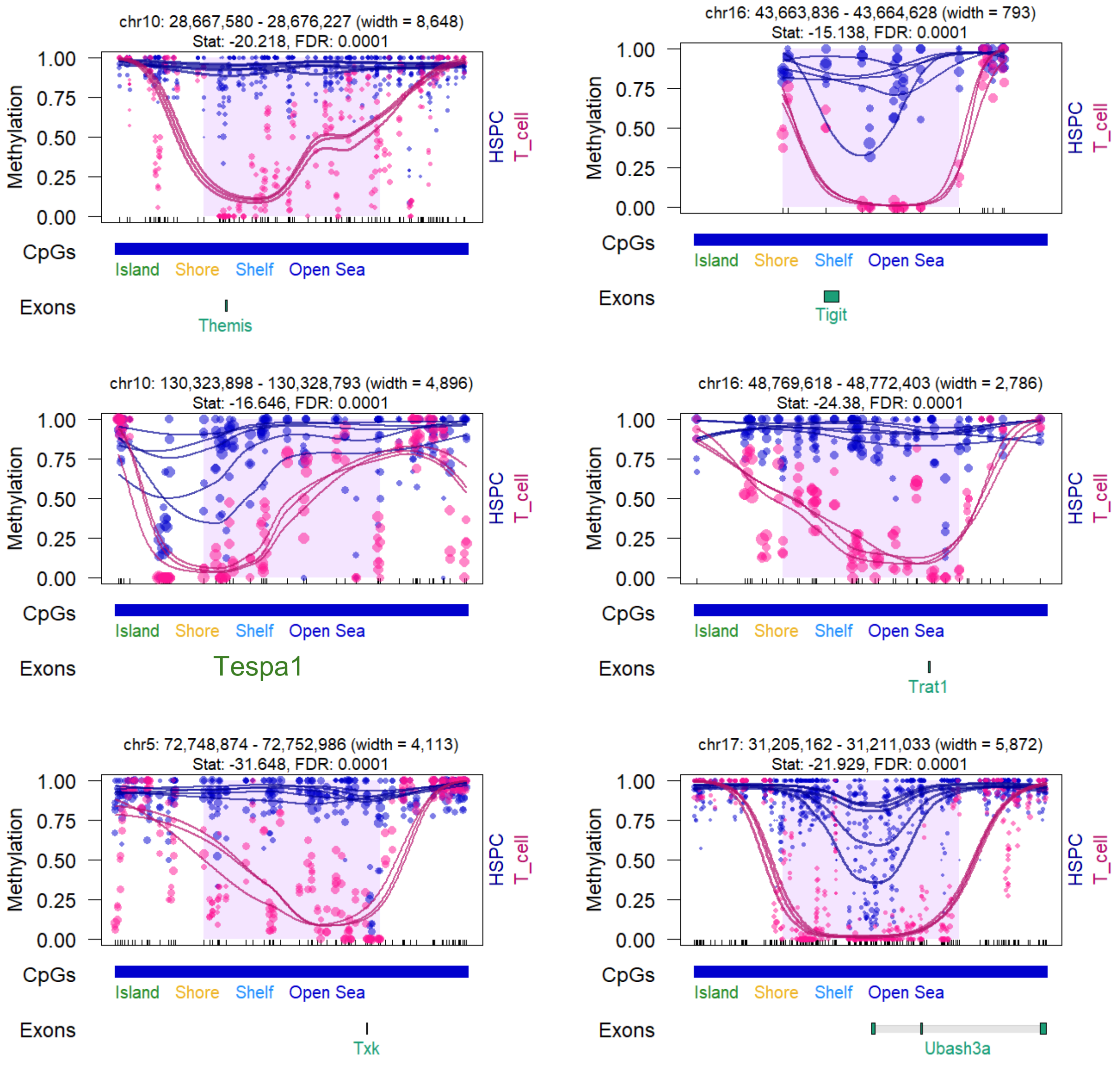
chr10: 28,667,580 - 28,676,227 (width = 8,648)
Stat: -20.218, FDR: 0.0001
Methylation
1.00
0.75
0.50
0.25
0.00
HSPC
T_cell
CpGs
Island Shore Shelf Open Sea
Exons
Themis
chr16: 43,663,836 - 43,664,628 (width = 793)
Stat: -15.138, FDR: 0.0001
Methylation
HSPC
T_cell
CpGs
Island Shore Shelf Open Sea
Exons
Tigit
chr10: 130,323,898 - 130,328,793 (width = 4,896)
Stat: -16.646, FDR: 0.0001
Methylation
HSPC
T_cell
CpGs
Island Shore Shelf Open Sea
Exons
Tespa1
chr16: 48,769,618 - 48,772,403 (width = 2,786)
Stat: -24.38, FDR: 0.0001
Methylation
HSPC
T_cell
CpGs
Island Shore Shelf Open Sea
Exons
Trat1
chr5: 72,748,874 - 72,752,986 (width = 4,113)
Stat: -31.648, FDR: 0.0001
Methylation
HSPC
T_cell
CpGs
Island Shore Shelf Open Sea
Exons
Txk
chr17: 31,205,162 - 31,211,033 (width = 5,872)
Stat: -21.929, FDR: 0.0001
Methylation
HSPC
T_cell
CpGs
Island Shore Shelf Open Sea
Exons
Ubash3a

### B cells (mm10)

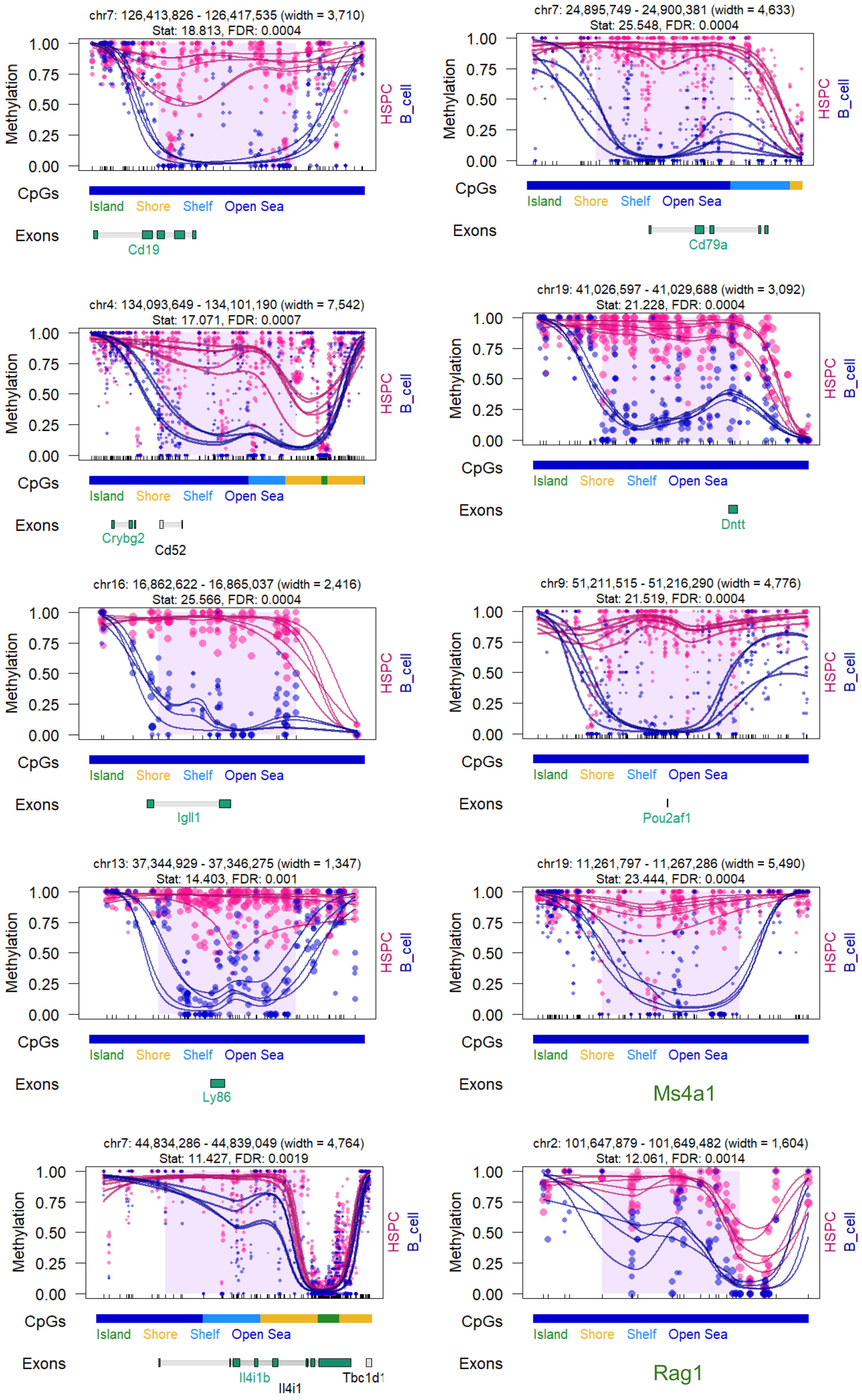

chr7: 126,413,826 - 126,417,535 (width = 3,710)
Stat: 18.813, FDR: 0.0004
Cd19
chr7: 24,895,749 - 24,900,381 (width = 4,633)
Stat: 25.548, FDR: 0.0004
Cd79a
chr4: 134,093,649 - 134,101,190 (width = 7,542)
Stat: 17.071, FDR: 0.0007
Crybg2
Cd52
chr19: 41,026,597 - 41,029,688 (width = 3,092)
Stat: 21.228, FDR: 0.0004
Dntt
chr16: 16,862,622 - 16,865,037 (width = 2,416)
Stat: 25.566, FDR: 0.0004
Igll1
chr9: 51,211,515 - 51,216,290 (width = 4,776)
Stat: 21.519, FDR: 0.0004
Pou2af1
chr13: 37,344,929 - 37,346,275 (width = 1,347)
Stat: 14.403, FDR: 0.001
Ly86
chr19: 11,261,797 - 11,267,286 (width = 5,490)
Stat: 23.444, FDR: 0.0004
Ms4a1
chr7: 44,834,286 - 44,839,049 (width = 4,764)
Stat: 11.427, FDR: 0.0019
Il4i1b
Il4i1
Tbc1d1
chr2: 101,647,879 - 101,649,482 (width = 1,604)
Stat: 12.061, FDR: 0.0014
Rag1
Methylation
1.00
0.75
0.50
0.25
0.00
HSPC
B_cell
CpGs
Island
Shore
Shelf
Open Sea
Exons

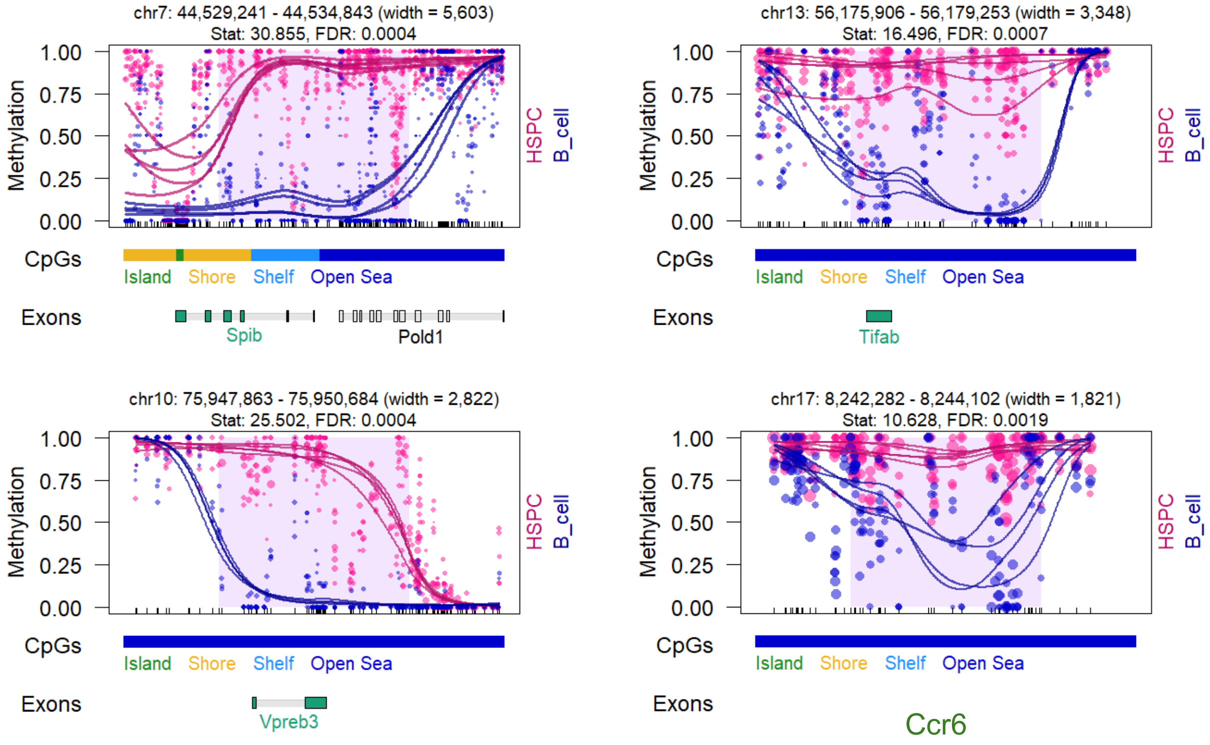

chr7: 44,529,241 - 44,534,843 (width = 5,603)
Stat: 30.855, FDR: 0.0004
Methylation
1.00
0.75
0.50
0.25
0.00
HSPC
B_cell
CpGs
Island
Shore
Shelf
Open Sea
Exons
Spib
Pold1
chr13: 56,175,906 - 56,179,253 (width = 3,348)
Stat: 16.496, FDR: 0.0007
Tifab
chr10: 75,947,863 - 75,950,684 (width = 2,822)
Stat: 25.502, FDR: 0.0004
Vpreb3
chr17: 8,242,282 - 8,244,102 (width = 1,821)
Stat: 10.628, FDR: 0.0019
Ccr6

#### monocytes / granulocytes (mm10)

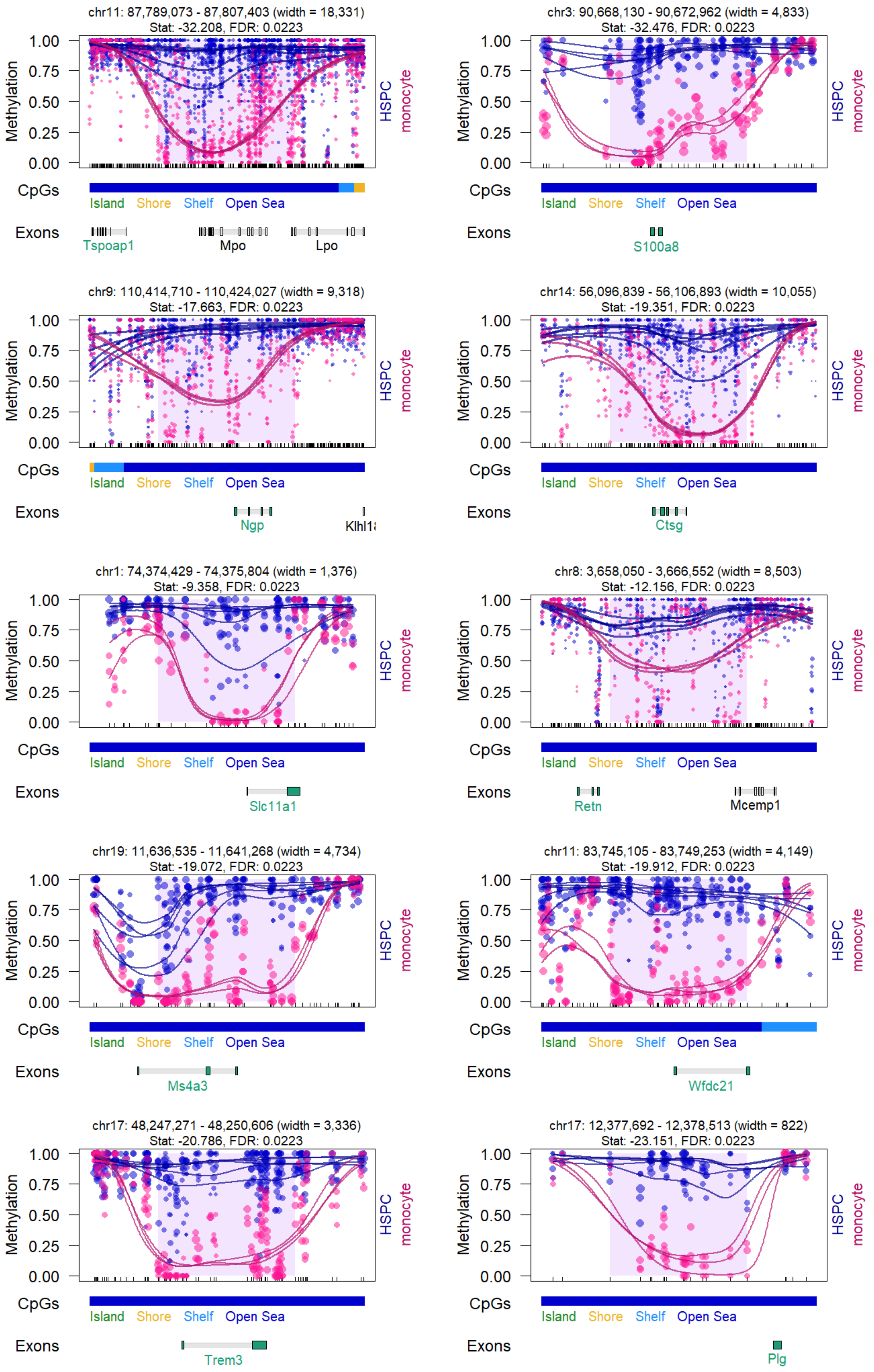

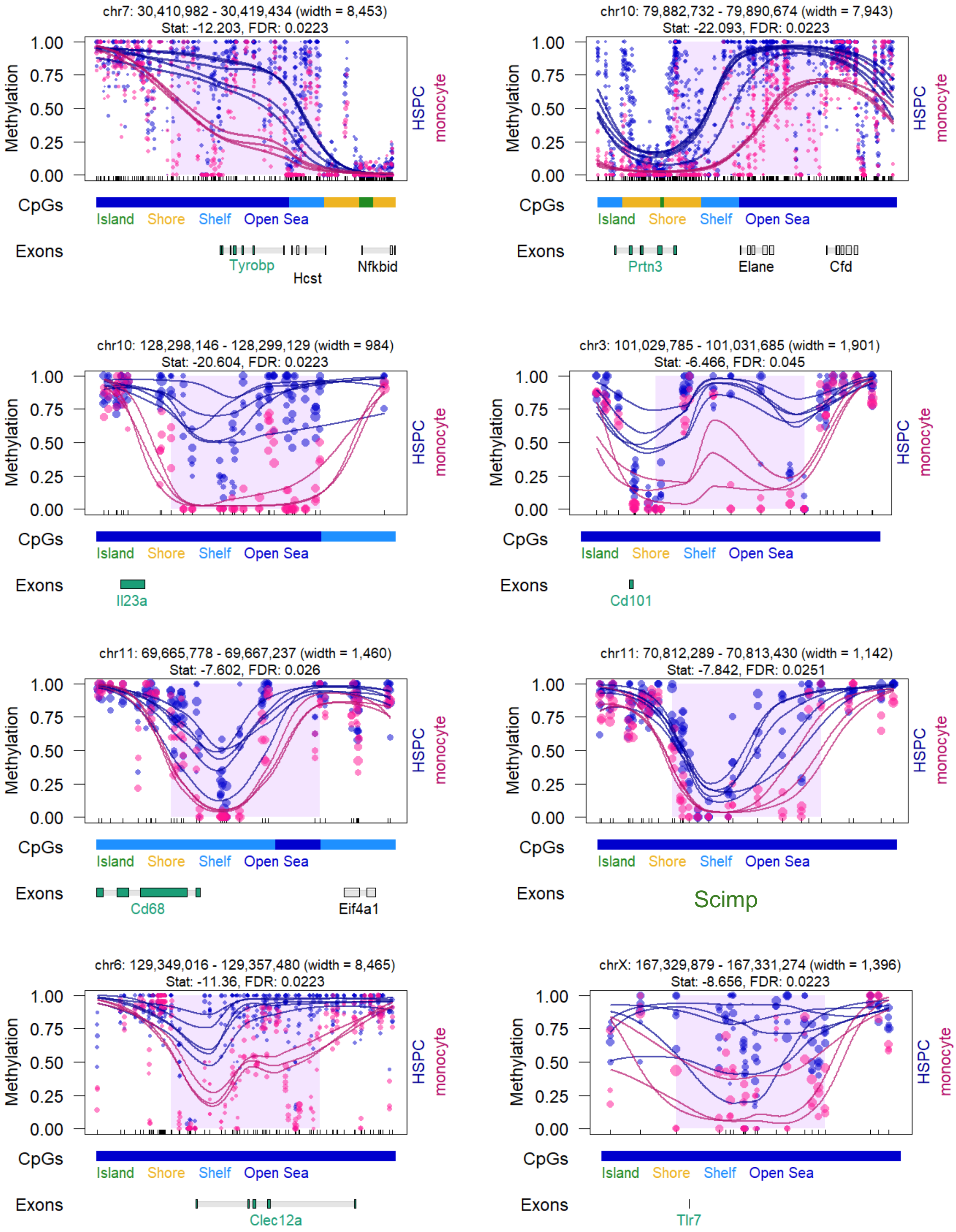
chr7: 30,410,982 - 30,419,434 (width = 8,453)
Stat: -12.203, FDR: 0.0223
Methylation
1.00
0.75
0.50
0.25
0.00
HSPC
monocyte
CpGs
Island
Shore
Shelf
Open Sea
Exons
Tyrobp
Hcst
Nfkbid
chr10: 79,882,732 - 79,890,674 (width = 7,943)
Stat: -22.093, FDR: 0.0223
Prtn3
Elane
Cfd
chr10: 128,298,146 - 128,299,129 (width = 984)
Stat: -20.604, FDR: 0.0223
Il23a
chr3: 101,029,785 - 101,031,685 (width = 1,901)
Stat: -6.466, FDR: 0.045
Cd101
chr11: 69,665,778 - 69,667,237 (width = 1,460)
Stat: -7.602, FDR: 0.026
Cd68
Eif4a1
chr11: 70,812,289 - 70,813,430 (width = 1,142)
Stat: -7.842, FDR: 0.0251
Scimp
chr6: 129,349,016 - 129,357,480 (width = 8,465)
Stat: -11.36, FDR: 0.0223
Clec12a
chrX: 167,329,879 - 167,331,274 (width = 1,396)
Stat: -8.656, FDR: 0.0223
Tlr7

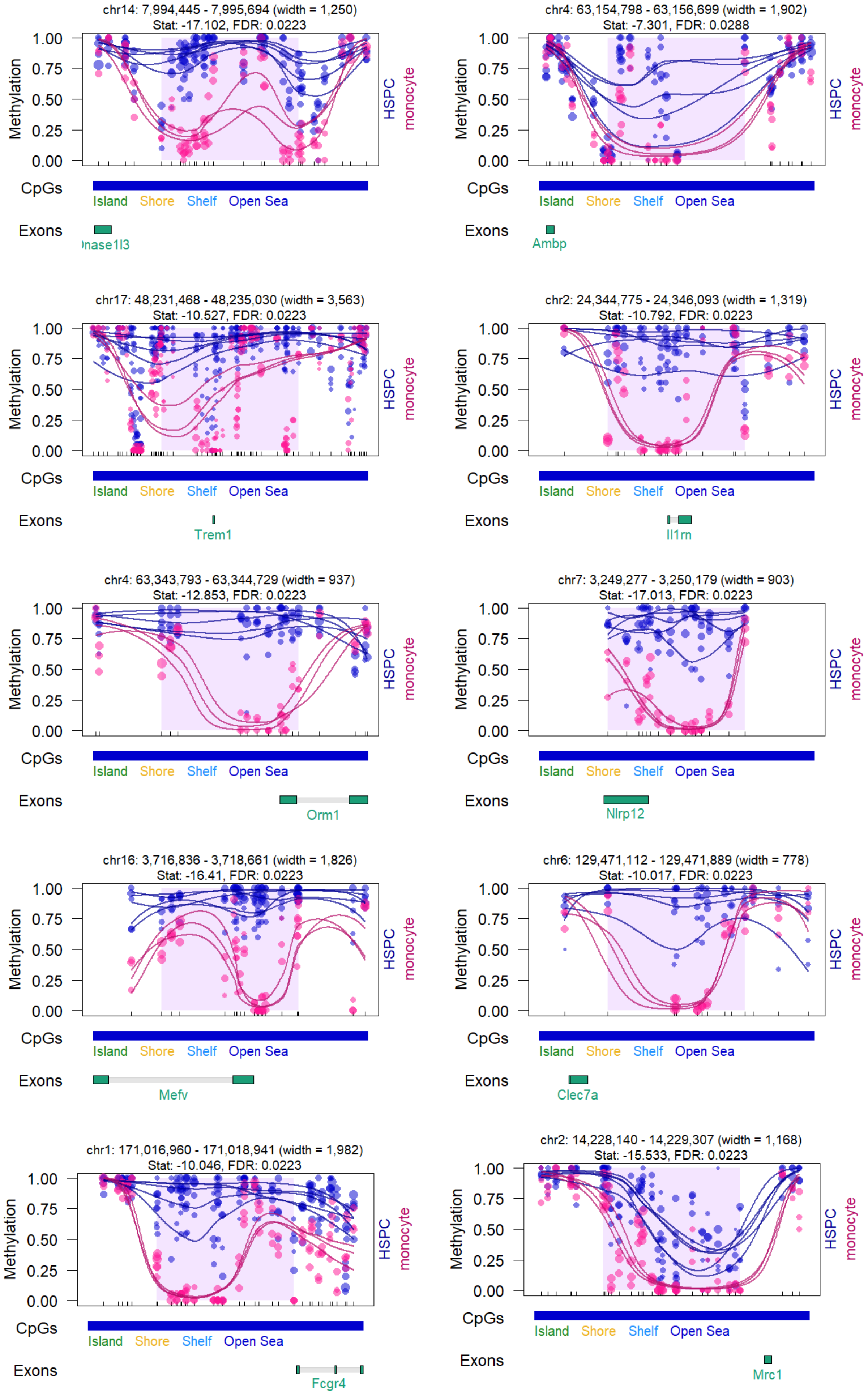
chr14: 7,994,445 - 7,995,694 (width = 1,250)
Stat: -17.102, FDR: 0.0223
nase1l3
chr4: 63,154,798 - 63,156,699 (width = 1,902)
Stat: -7.301, FDR: 0.0288
Ambp
chr17: 48,231,468 - 48,235,030 (width = 3,563)
Stat: -10.527, FDR: 0.0223
Trem1
chr2: 24,344,775 - 24,346,093 (width = 1,319)
Stat: -10.792, FDR: 0.0223
Il1rn
chr4: 63,343,793 - 63,344,729 (width = 937)
Stat: -12.853, FDR: 0.0223
Orm1
chr7: 3,249,277 - 3,250,179 (width = 903)
Stat: -17.013, FDR: 0.0223
Nlrp12
chr16: 3,716,836 - 3,718,661 (width = 1,826)
Stat: -16.41, FDR: 0.0223
Mefv
chr6: 129,471,112 - 129,471,889 (width = 778)
Stat: -10.017, FDR: 0.0223
Clec7a
chr1: 171,016,960 - 171,018,941 (width = 1,982)
Stat: -10.046, FDR: 0.0223
Fcgr4
chr2: 14,228,140 - 14,229,307 (width = 1,168)
Stat: -15.533, FDR: 0.0223
Mrc1
Methylation
1.00
0.75
0.50
0.25
0.00
HSPC
monocyte
CpGs
Island
Shore
Shelf
Open Sea
Exons
