## Supplementary figures and images for "The TET-dependent DNA demethylation pathway is the driving force of hematopoiesis"

### Supplementary Data S6

#### additional genes

##### T cells (hg38)

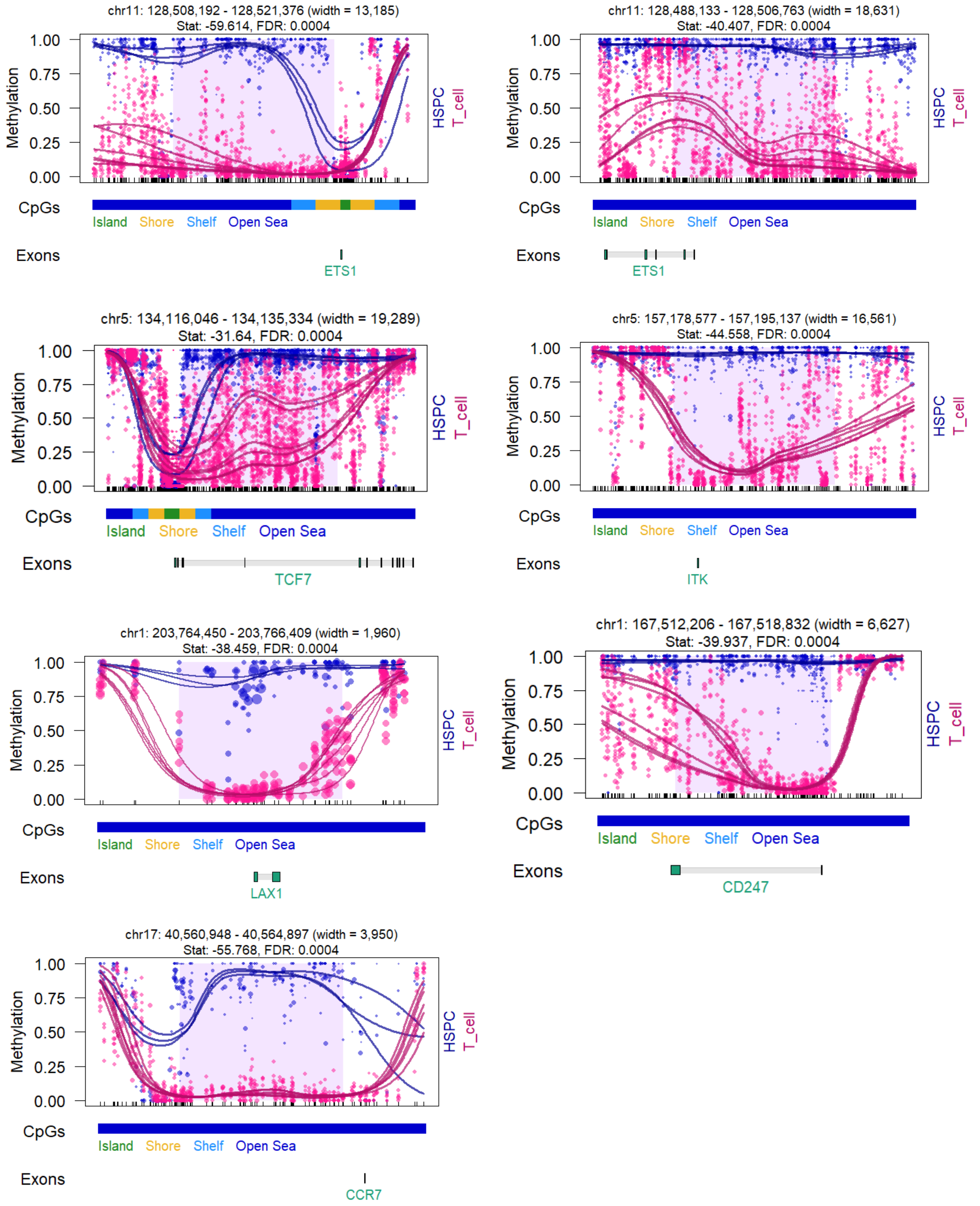

#### B cells (hg38)

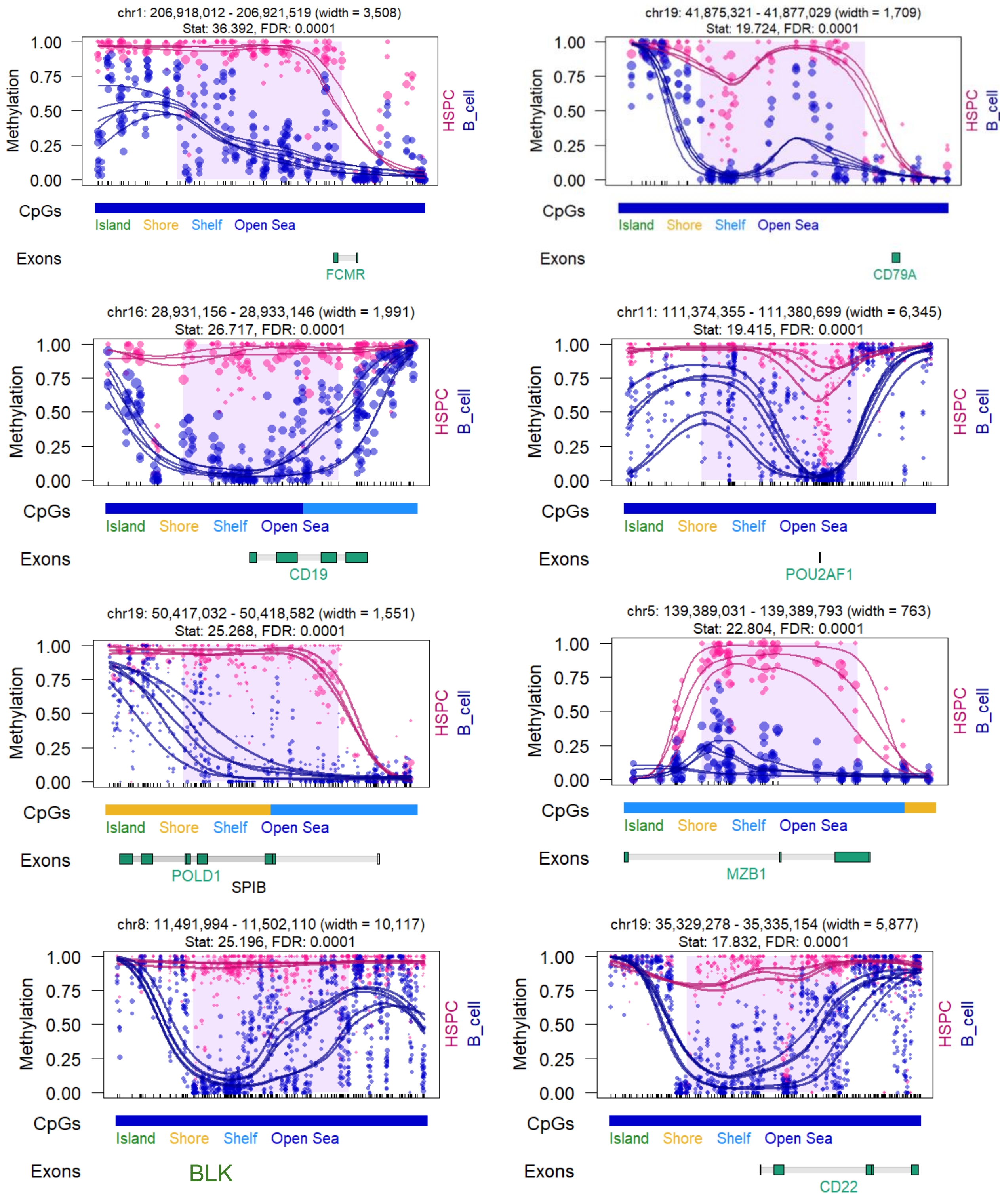


#### Monocytes (hg38)

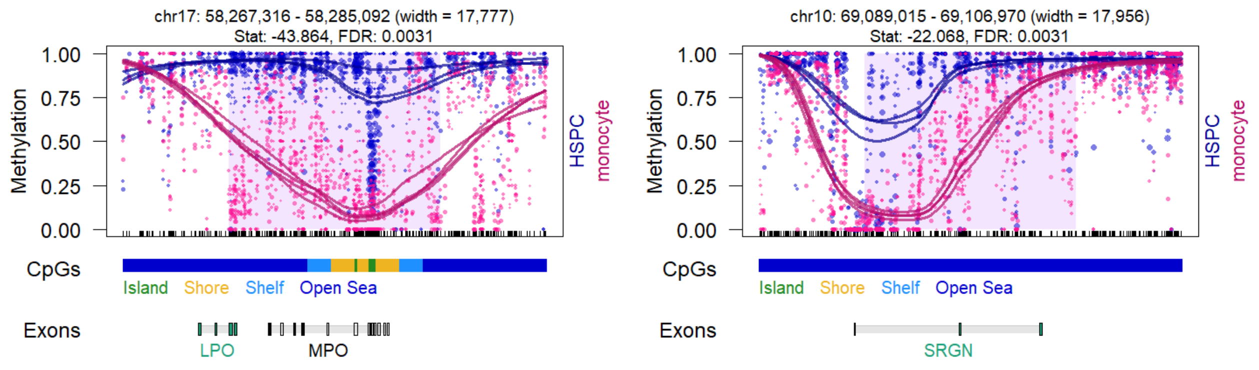

#### all genes

##### T cells (hg38)

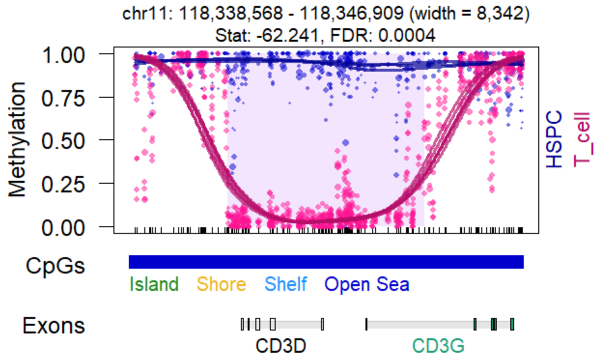


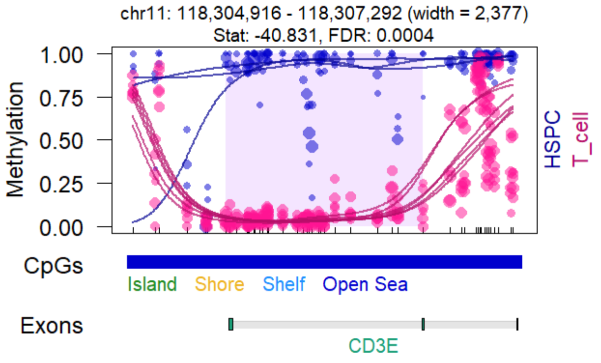


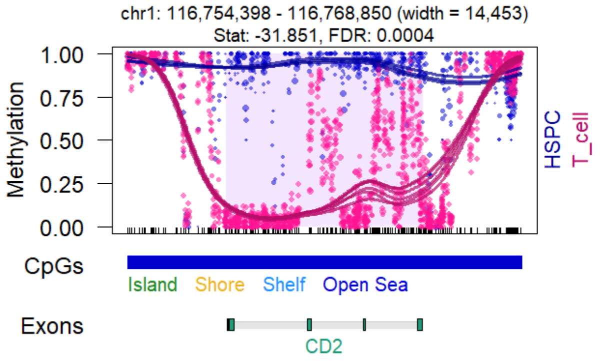


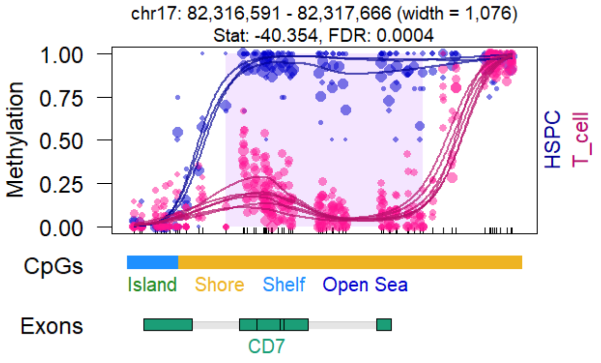


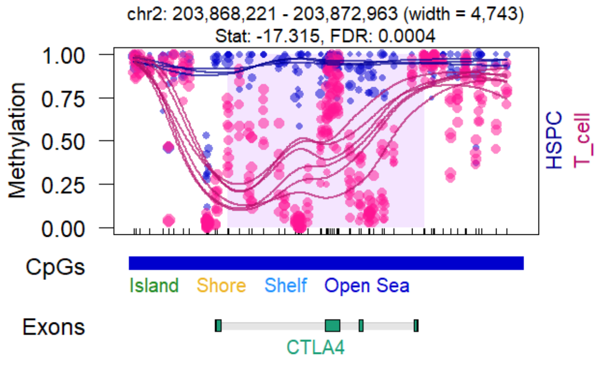


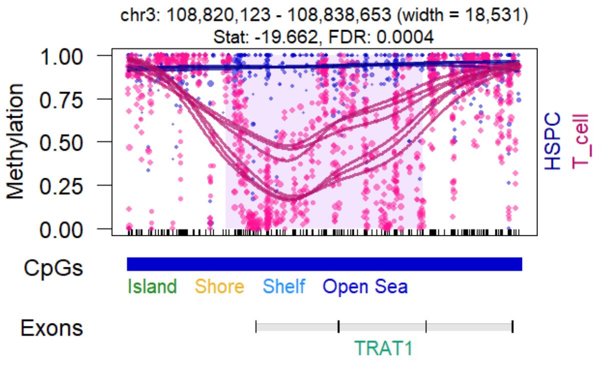


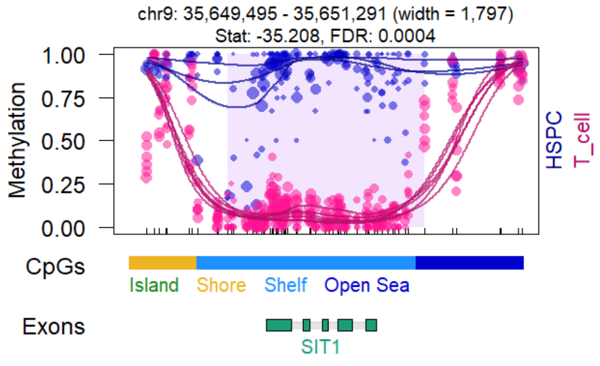


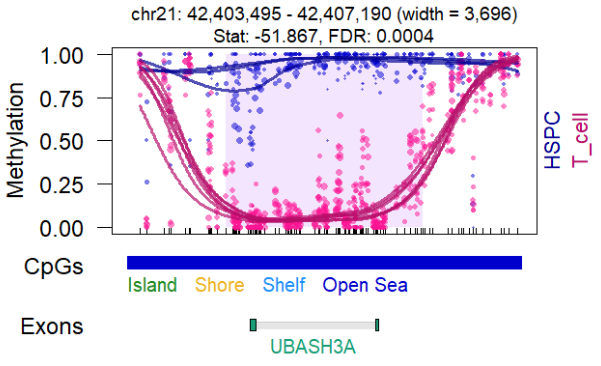

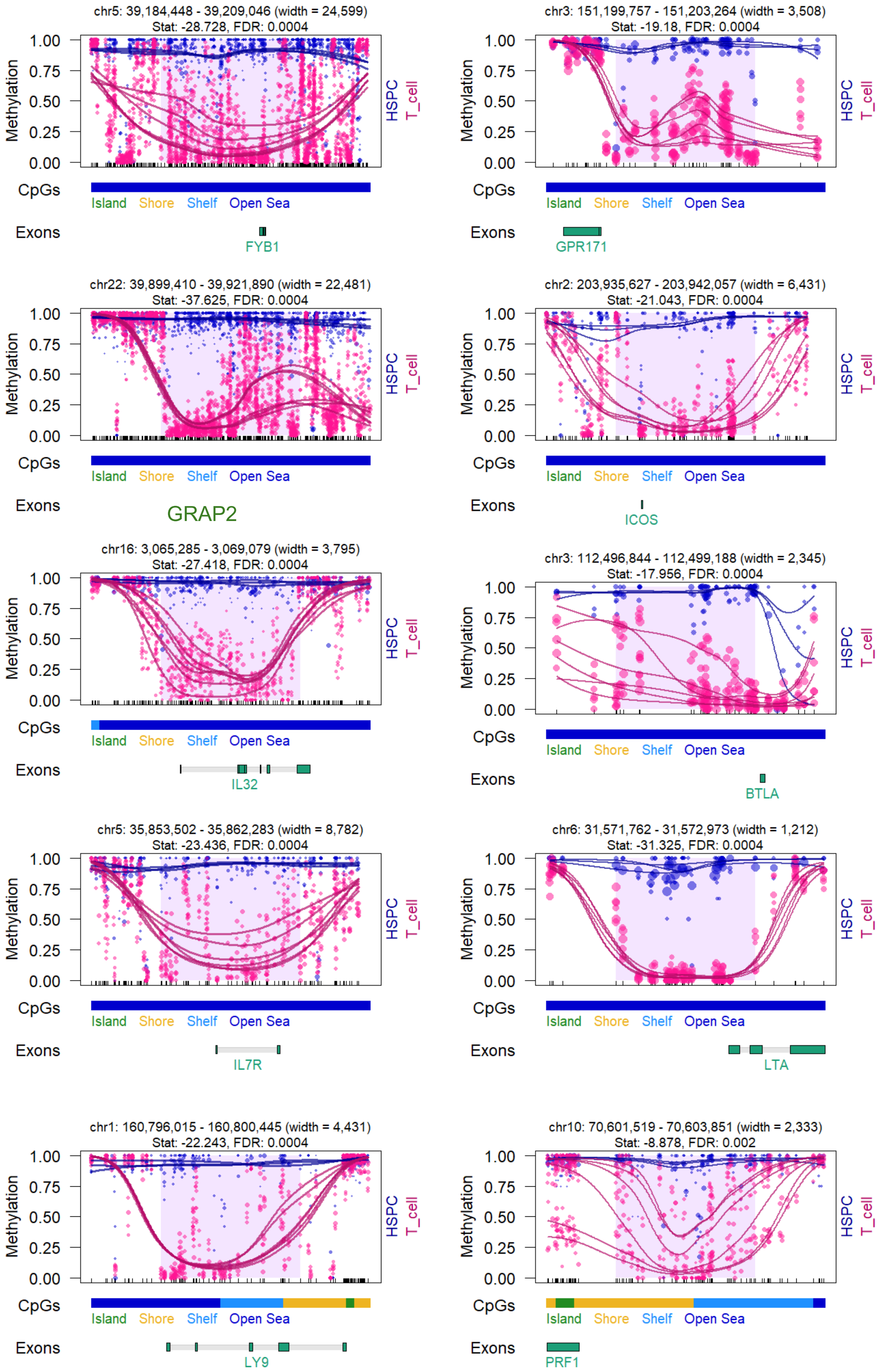

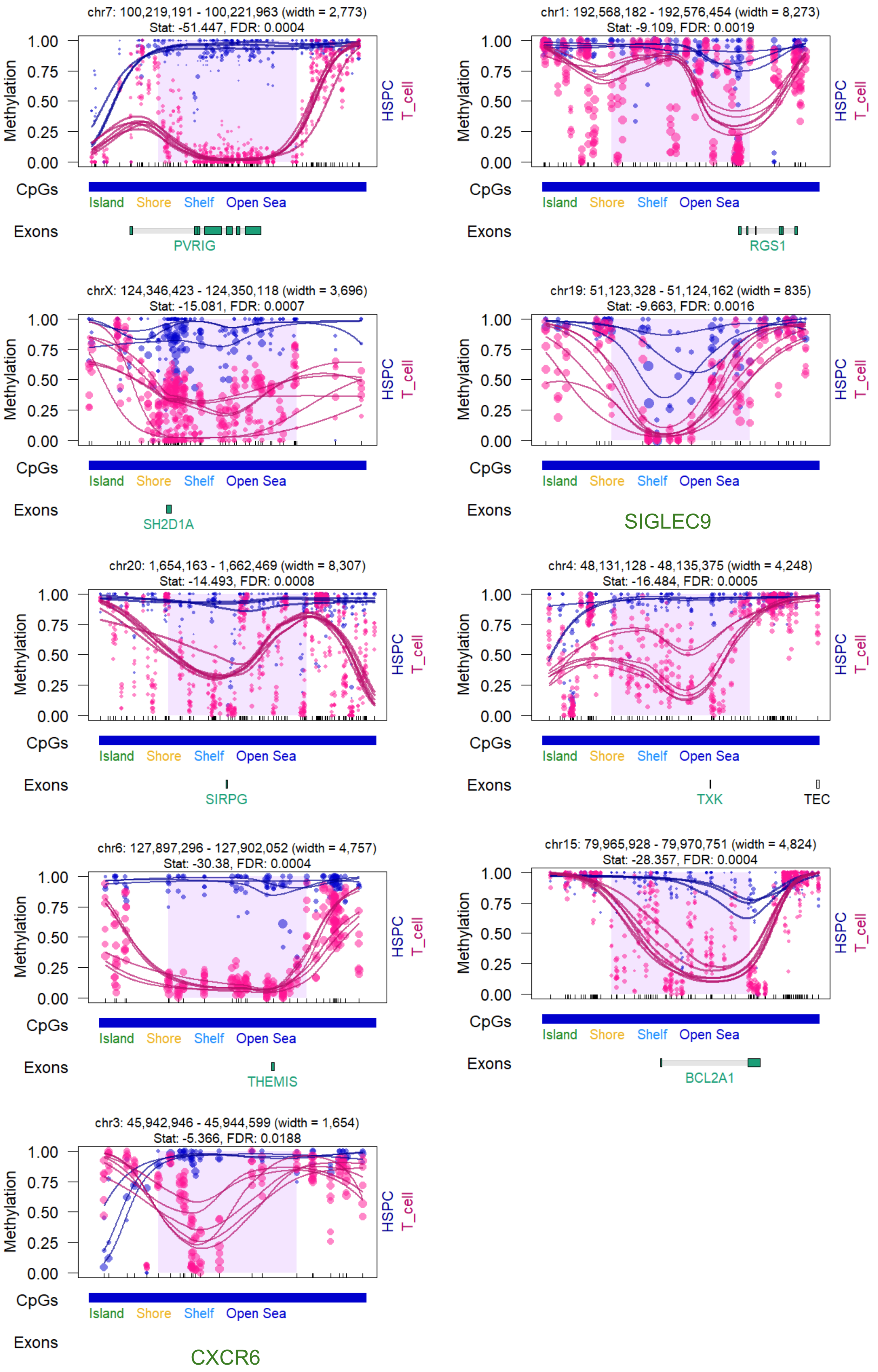

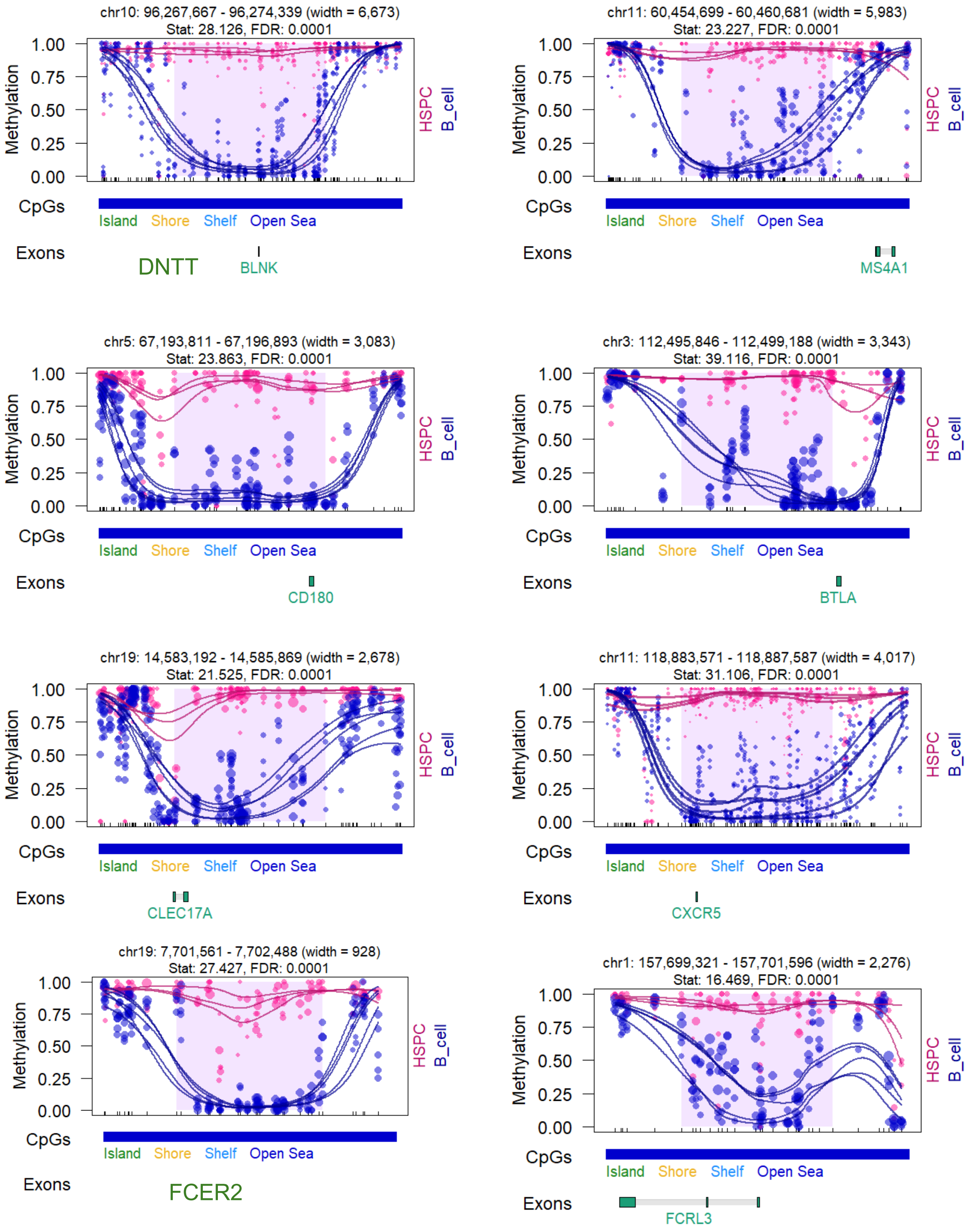

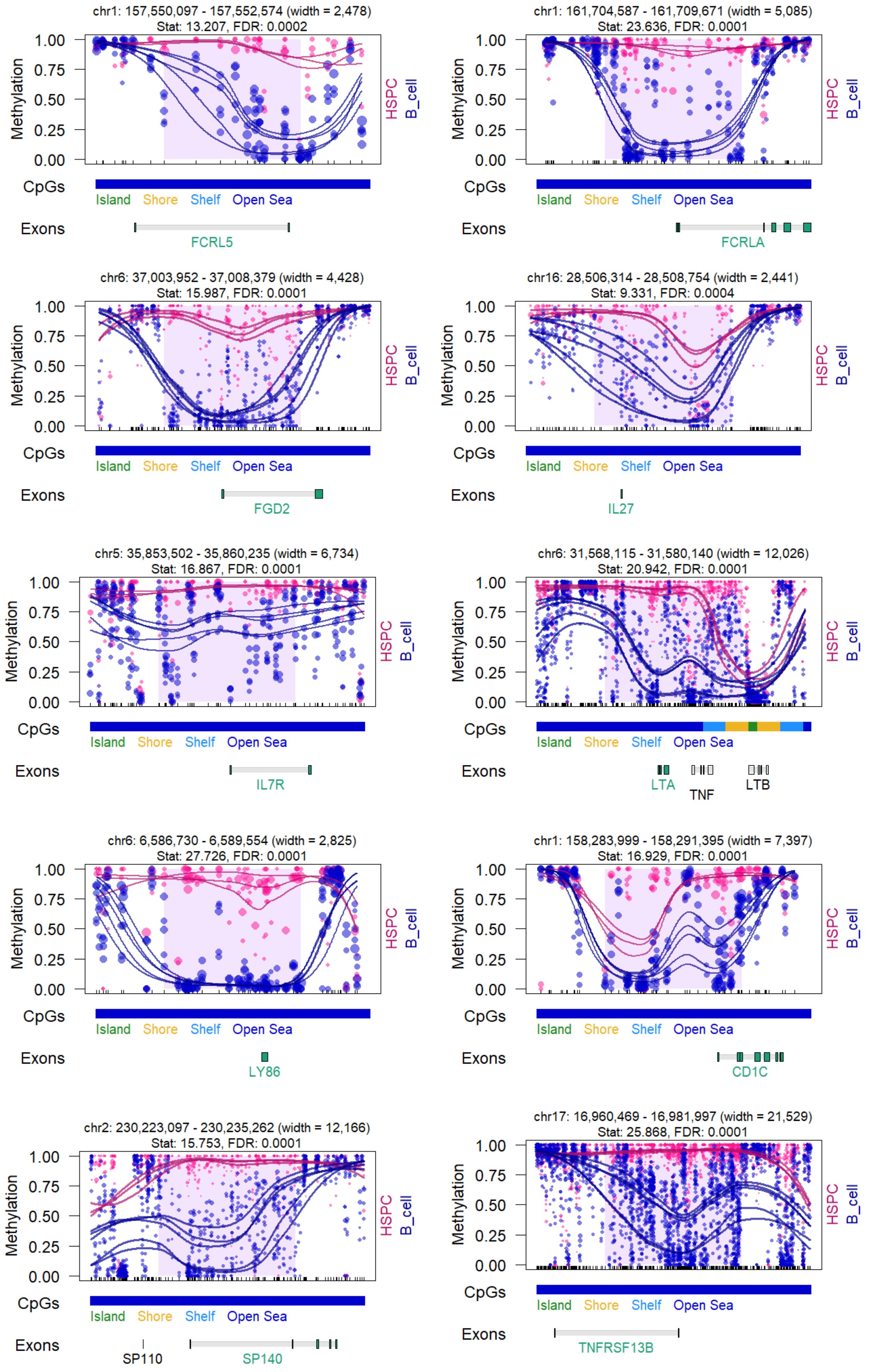

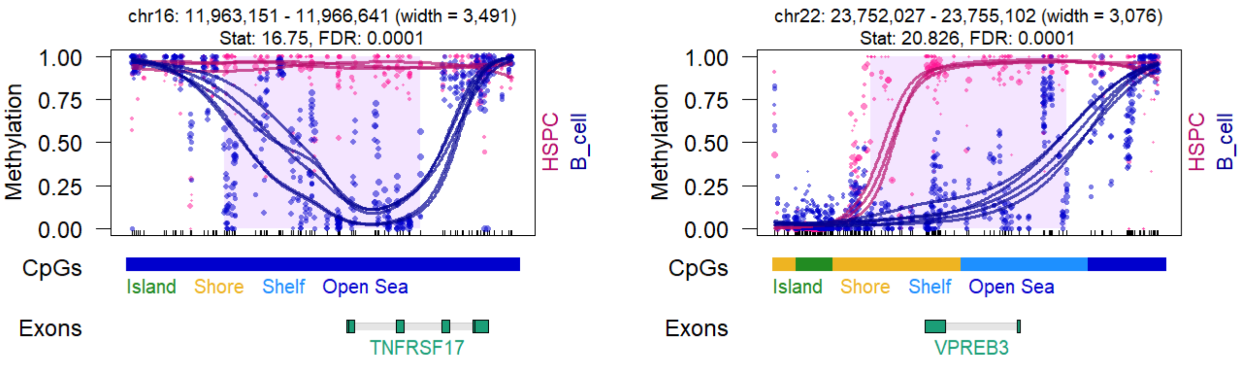


#### Monocytes (hg38)

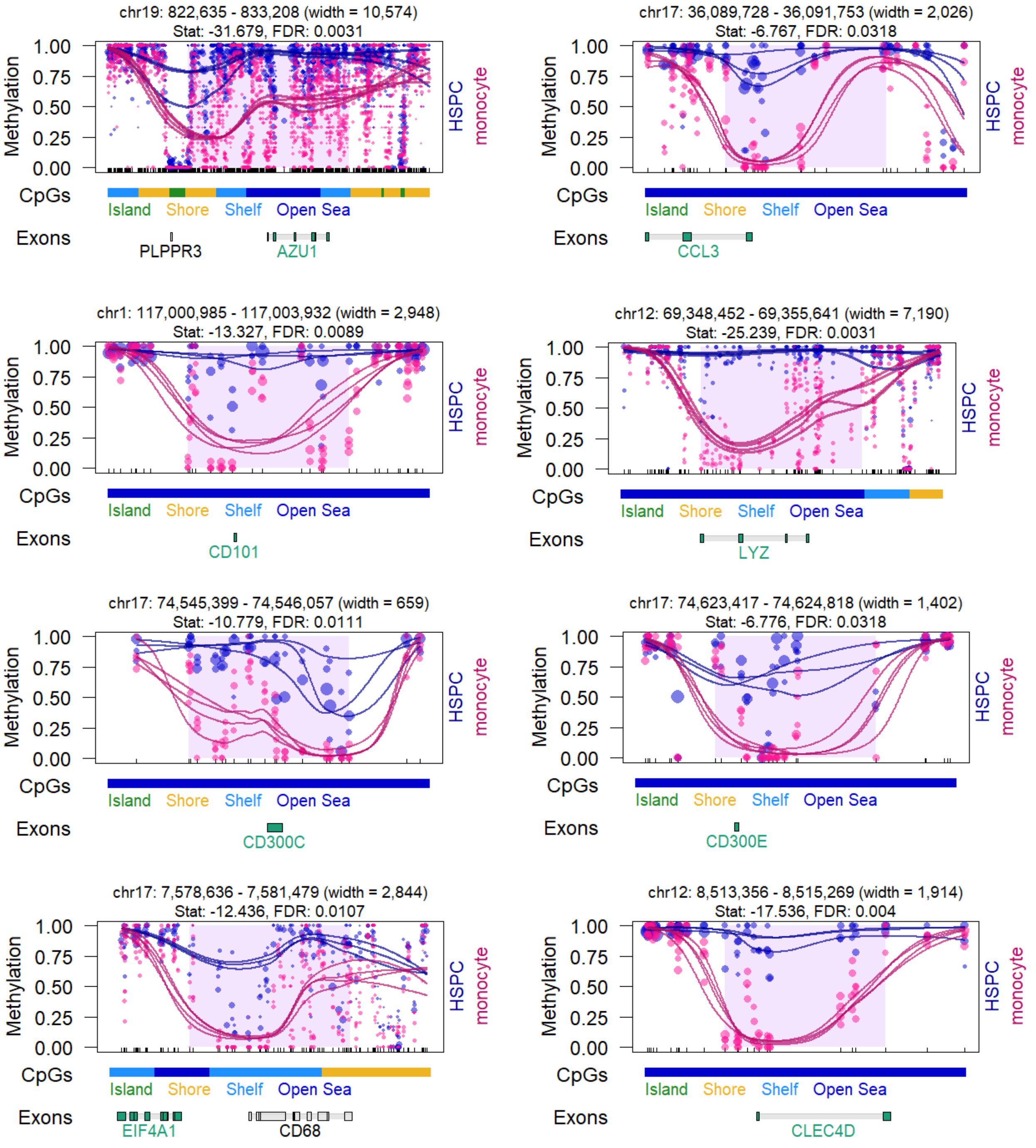

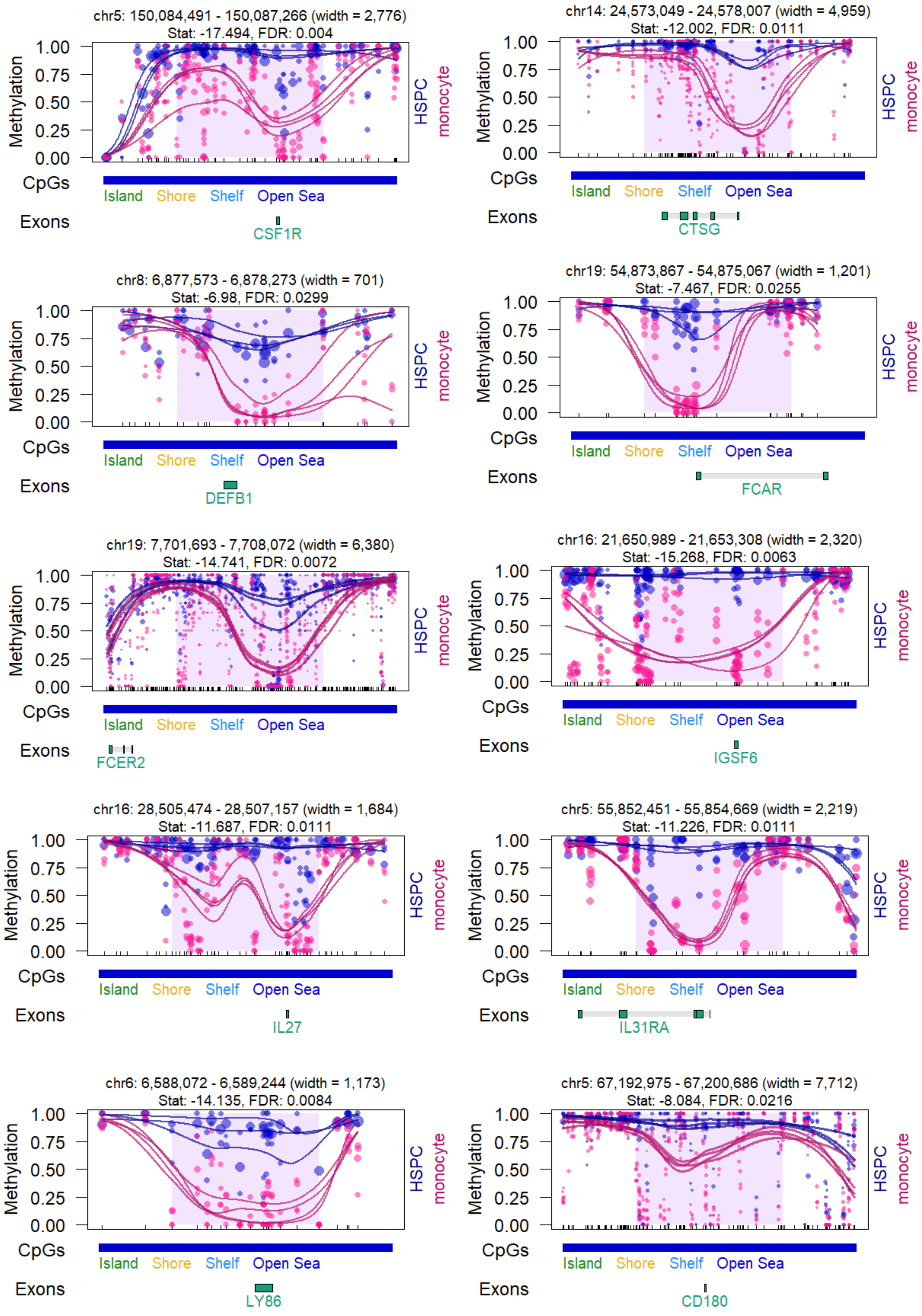

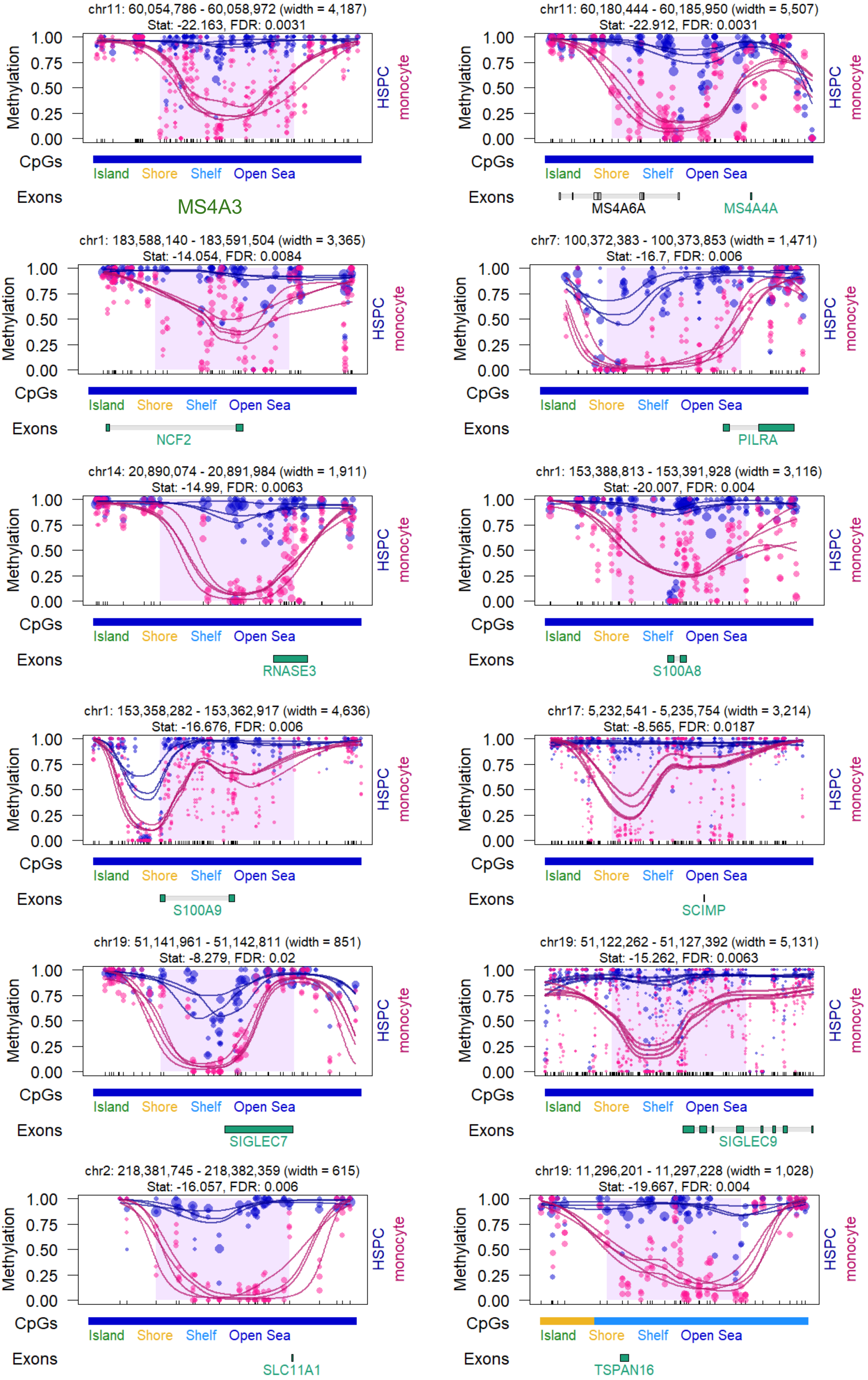
